# Targeted Transcriptional Activation using a CRISPR-Associated Transposon System

**DOI:** 10.1101/2023.09.12.557412

**Authors:** Andrea M. Garza Elizondo, James Chappell

## Abstract

Synthetic perturbation of gene expression is central to our ability to reliably uncover genotype-phenotype relationships in microbes. Here, we present a novel gene transcription activation strategy that uses the *Vibrio cholerae* CRISPR-Associated Transposon (CAST) system to selectively insert promoter elements upstream of genes of interest. Through this strategy, we show robust activation of both recombinant and endogenous genes across the *E. coli* chromosome. We then demonstrate precise tuning of expression levels by exchanging the promoter elements being inserted. Finally, we demonstrate that CAST activation can be used to synthetically induce ampicillin-resistant phenotypes in *E. coli*.

## Introduction

Understanding the relationships between genotype and phenotype is one of the grand challenges of biology that has driven the development of molecular strategies for manipulating and deciphering gene function and connectivity. A mainstay approach to studying gene function is the perturbation of endogenous gene expression followed by the observation of resulting phenotypic changes. Perturbations meant to activate transcription or overexpress a gene of interest are known as knock-up studies and their inactivation counterparts as knock-out or knock-down studies. Knock-up studies have been conducted through a variety of methods, including transposon mutagenesis, plasmid-based overexpression, and insertion of promoters through recombination. Through these studies, genes have been identified that confer resistance or tolerance to antibiotics,^1–5^ oxidizing agents,^6^ metals,^7^ and other toxins.^2,8^ Likewise, alternate enzymatic activities^9^ and the effects of overexpression on growth^10^ have been identified through knock-up experiments. In eukaryotic cells, knock-up studies have also been used to uncover the organization and connectivity in regulatory networks.^11^ Overall, the activation and overexpression of genes has proven to be a fruitful mechanism by which to study genes.

Recently, CRISPR-based strategies for gene activation have been developed, expanding our ability to achieve targeted overexpression of genes of interest. There are two main strategies for activating transcription in prokaryotes using CRISPR-based editors and regulators. One strategy uses CRISPR-Cas9 DNA editing for the insertion of promoter elements upstream of target genes, also known as promoter knock-in.^12–14^ CRISPR-Cas9 knock-in allows for precise, scar-less insertion of promoters. The other strategy, CRISPR activation (CRISPRa), uses a catalytically dead Cas9 (dCas9) to localize activation domains upstream of a target gene’s promoter, thereby prompting that gene’s expression.^15–19^ The same dCas9 used for activation can be repurposed for CRISPR interference (CRISPRi), which makes this strategy attractive for large-scale screens due to the ability to create libraries of single guide RNAs (sgRNAs) for both CRISPRi and CRISPRa in a single experiment.^16,17,19–21^ While both strategies confer advantages unique to their systems, challenges remain for applying them at a scale (e.g. for genome-wide studies). For instance, promoter knock-in by CRISPR-Cas requires a customized DNA donor with large homologous arms specific to each edit site (e.g., ∼400 bp).^22^ This can increase the complexity of library design as both the sgRNA and the DNA donor must be varied simultaneously. On the other hand, CRISPRa benefits from simplified library design, requiring only a sgRNA for each targeted gene. However, different sets of challenges remain (as reviewed in ^23^) that include periodical activation patterns which make it difficult to activate diverse promoters,^16,18,24,25^ and the use of activation domains that can require customization for different promoter types^24,26^ and hosts.^18,24^

Here, we contribute a novel transcription activation strategy that complements existing CRISPR-based systems, using a recently discovered CRISPR-Associated Transposon (CAST). The *Vibrio cholerae* CAST system is a naturally occurring hybrid of type I-F CRISPR-Cas systems and *E. coli* Tn7-like transposons that has retained the guide RNA-processing and RNA-guided binding properties of CRISPR-Cas systems alongside the transposition mechanism of Tn7.^27,28^ The result is the targeted insertion of DNA transposons downstream of sites complementary to the encoded CRISPR RNA (crRNA). This unique mechanism of RNA-guided transposition provides the CAST system with several attractive attributes from a gene perturbation standpoint. For one, the CAST system has already shown great versatility for genome knock-downs^28^ and knock-outs in multiple species for creation of auxotrophic strains,^29^ for metabolic pathway engineering,^30^ and for microbiome programming.^31^ Furthermore, the CAST system can leverage universal transposons, analogous to the DNA donor used in CRISPR-Cas9 editing, that are interchangeable between different target sites. This makes the system amenable for library strategies wherein one transposon cargo can be used with different crRNAs and target sites.^29,30,32,33^ While CAST has only been used for repression, we reasoned that by encoding outward-facing promoter elements within the transposon cargo and inserting these cargos upstream of silent or weakly-expressed genes, we could adapt the CAST system to also function for activation.

In this paper, we explore the activation potential of the CAST systems. To demonstrate this novel function, we first use the CAST system to insert a strong constitutive promoter as a transposon cargo upstream of fluorescent reporter genes, leading to robust transcription of the fluorescent gene of interest. We then demonstrate the fine-tuning of transcriptional activation using variable strength and inducible promoter systems as transposon cargos. Furthermore, we demonstrate the utility of this approach in non-synthetic contexts by activating the transcription of endogenous *E. coli* genes. Finally, we demonstrate that CAST can be used to identify endogenous genes that confer antibiotic resistance when overexpressed in *E. coli*.

## Results

### Engineering the CAST system for efficient transcriptional activation

As a starting point, our first goal was to ascertain if CAST can be used to activate the expression of genome-encoded genes through the transposition of outward-facing promoter elements upstream of a targeted gene of interest (GOI) (**Figure 1A**). To test this, we focused on adapting the CAST system derived from the *Vibrio cholerae* Tn6677 transposon, which is composed of a crRNA-encoding CRISPR array, seven proteins, and a transposon containing a DNA cargo flanked by transposon left end (LE) and right end (RE) elements.^33^ Prior work has shown that this CAST system can be targeted via the crRNA, resulting in high-efficiency insertion of the transposon ∼49 bp downstream of the crRNA binding site.^27,33,34^ Interestingly, while the transposon can be inserted in either direction (*i*.*e*., RE-LE or LE-RE orientation) it shows a significant bias for RE-LE insertions.^27,33,34^ Given these features, we hypothesized that the transposition of promoter elements as the transposon cargo could serve as a reliable strategy for activating gene expression (**Figure 1A**).

**Figure 1.**
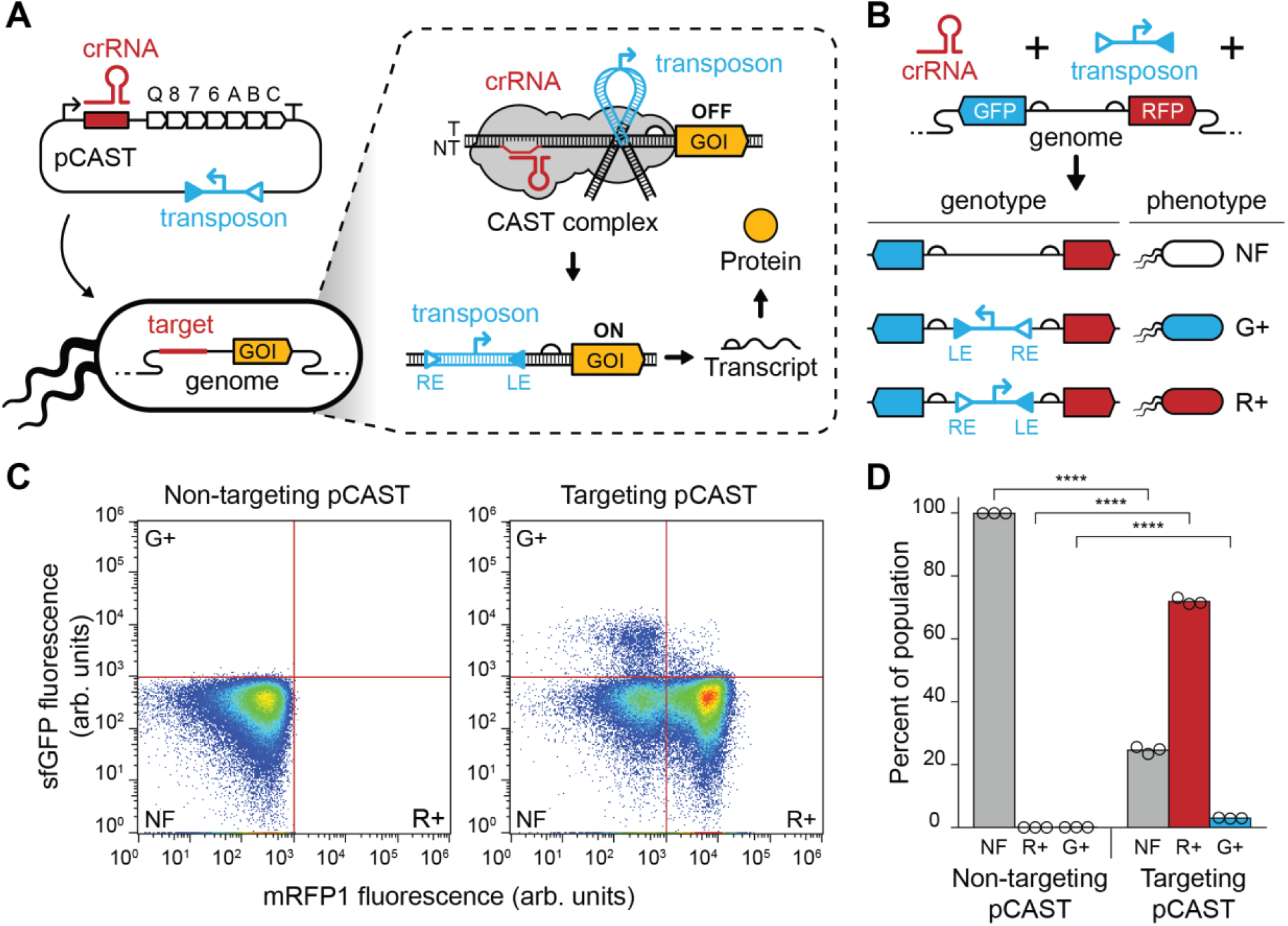
CAST can activate transcription through the insertion of transcriptional promoters. (**A**) Schematic of the mechanism of transcriptional activation by CAST editing through promoter insertion. The pCAST plasmid encodes an operon with the crRNA, the CAST complex, and the transposon with a promoter cargo and the left end (LE) and right end (RE) elements. The crRNA is designed to target the CAST complex upstream of a gene of interest (GOI). Insertion of the transposon in the right-to-left end (RE-LE) direction results in transcriptional activation of the GOI. (**B**) Schematic of the engineered reporter strain and outcomes of CAST editing. An *E. coli* strain was engineered to contain sfGFP (GFP) and mRFP1 (RFP) reporters lacking promoter elements. CAST editing with a transposon containing a promoter cargo results in cells with mRFP1 fluorescence (R+) if the transposon is inserted in the right-to-left orientation, sfGFP fluorescence (G+) if the transposon is inserted in the left-to-right orientation, and no fluorescence (NF) if editing does not occur. (**C**) Single-cell fluorescence analysis of *E. coli* cells edited with CAST. Flow cytometry data showing the single-cell sfGFP and mRFP1 fluorescence of *E. coli* cells transformed with a non-targeting (left panel) and targeting (right panel) pCAST plasmid. Red lines indicate thresholds used to determine NF, G+, and R+ cell populations. Data are representatives of *n* = 3, with replicates shown in **Supplementary Fig. 1**. (**D**) CAST editing outcomes of *E. coli* cells transformed with a non-targeting and targeting pCAST plasmid. Populations are labeled as in (B), bars represent averages of *n* = 3 replicates. Statistical significance between populations is indicated by asterisks (Student’s T-Test, p < 0.0005).

To test this, we first created a reporter *E. coli* strain in which fluorescent sfGFP and mRFP1 lacking transcriptional promoters were encoded within the genome in divergent directions (**Figure 1B**). We reasoned we could use this strain to measure the overall efficiency of editing and directionality of insertion by designing a plasmid construct (called pCAST) containing a CAST system, a transposon cargo containing a strong constitutive promoter facing the LE, and a crRNA to target the intergenic region between the two reporter genes. The crRNA was designed to target the template strand upstream of mRFP1, which we presumed would result in predominantly RE-LE insertions and transcription activation of mRFP1. Although transposon insertion directionality is biased toward the RE-LE direction, we also expected low frequency of insertion in the LE-RE direction, resulting in sfGFP fluorescence, and low frequency of no insertion, resulting in no fluorescence (**Figure 1B**). The pCAST plasmid was transformed into the sfGFP-mRFP1 reporter strain followed by an incubation period of 24 hr at 30°C in liquid culture, a method previously shown to result in high transposition efficiencies.^33^ Fluorescence was then quantified using single-cell flow cytometry analysis. The fluorescence of the post-incubation bacterial population showed both high transposition and transcription activation of mRFP1 (**Figure 1C, Supplementary Figure 1**). Specifically, the fraction of the population with observed fluorescence was 74.8%, 96.1% of which were from mRFP1 positive cells for a final mRFP1 activation efficiency of 71.9% (**Figure 1D**). We observed a relatively low frequency of no insertion events (24.6%) and a significantly lower rate of LE-RE insertions (2.95%) in agreement with the prior studies.^33^

Next, we sought to determine the distance between the crRNA binding site and the transposon insertion in transcriptionally activated mRFP1 cells. To do this, we isolated mRFP1 positive cells using fluorescence-activated cell sorting (FACS) and performed next-generation sequencing of the sfGFP-mRFP1 genomic locus (**Supplementary Figure 2**). This analysis revealed that the majority (68.4%) of insertions were located 48 bp downstream of the 3’ end of the crRNA binding site, in agreement with insertion distances observed in prior literature.^33,34^ Additionally, 26.6% of the remaining insertions occurred at low frequency within a 5 bp window centered on 48 bp.

Having confirmed that our CAST activation strategy yielded high levels of transcriptional activation, we were interested in determining possible effects on long-term activation that resulted from the binding of the CAST proteins. Specifically, prior work has shown that binding by the CAST complex can have repressive effects on nearby genes,^28^ possibly affecting the activation achieved through our strategy if the CAST complex remains in the cell. Likewise, we reasoned that the binding of the TnsA and TnsB proteins to the transposon ends^28^ could be blocking polymerase read-through into the gene of interest, reducing activation levels. To investigate this, we transformed an empty vector with no CAST complex, an mRFP1 targeting pCAST plasmid, and a non-targeting pCAST plasmid into sfGFP-mRFP1 reporter strains already containing a RE-LE-oriented transposon (**Supplementary Figure 3**). Fluorescence measurement showed modest reductions in mRFP1 expression levels only in the presence of a non-targeting pCAST, suggesting that the presence of the CAST complex or TnsB was not dramatically reducing transcription readthrough.

### Using the CAST system to achieve tunable transcription activation

After demonstrating that CAST can be used to achieve transcription activation, we next sought to tune the level of activation. To do so, we exchanged the promoter encoded in the transposon cargo for different strength constitutive promoter elements and performed CAST editing to activate mRFP1 expression. As expected, the fluorescence intensity of the mRFP1 positive cells decreased as the strength of the promoter decreased (**Figure 2A, Supplementary Figure 4**). An alternative strategy to achieve tunable control of gene activation is with inducible promoters. To test this, we exchanged the promoter within the transposon for the anhydrotetracycline (aTc)-inducible promoter (PTet) and its cognate repressor, *tetR*. This plasmid was then transformed into the dual reporter strain, and a population containing the transposon inserted in the RE-LE orientation was isolated. Induction of the promoter with increasing concentrations of aTc showed increasing mRFP1 fluorescence intensity, showcasing the ability to tune the level of activation achieved by the promoter (**Figure 2B, Supplementary Figure 5**). Taken together, these results show the potential use of the CAST system as a targeted and tunable transcription activator via the insertion of promoter elements.

**Figure 2.**
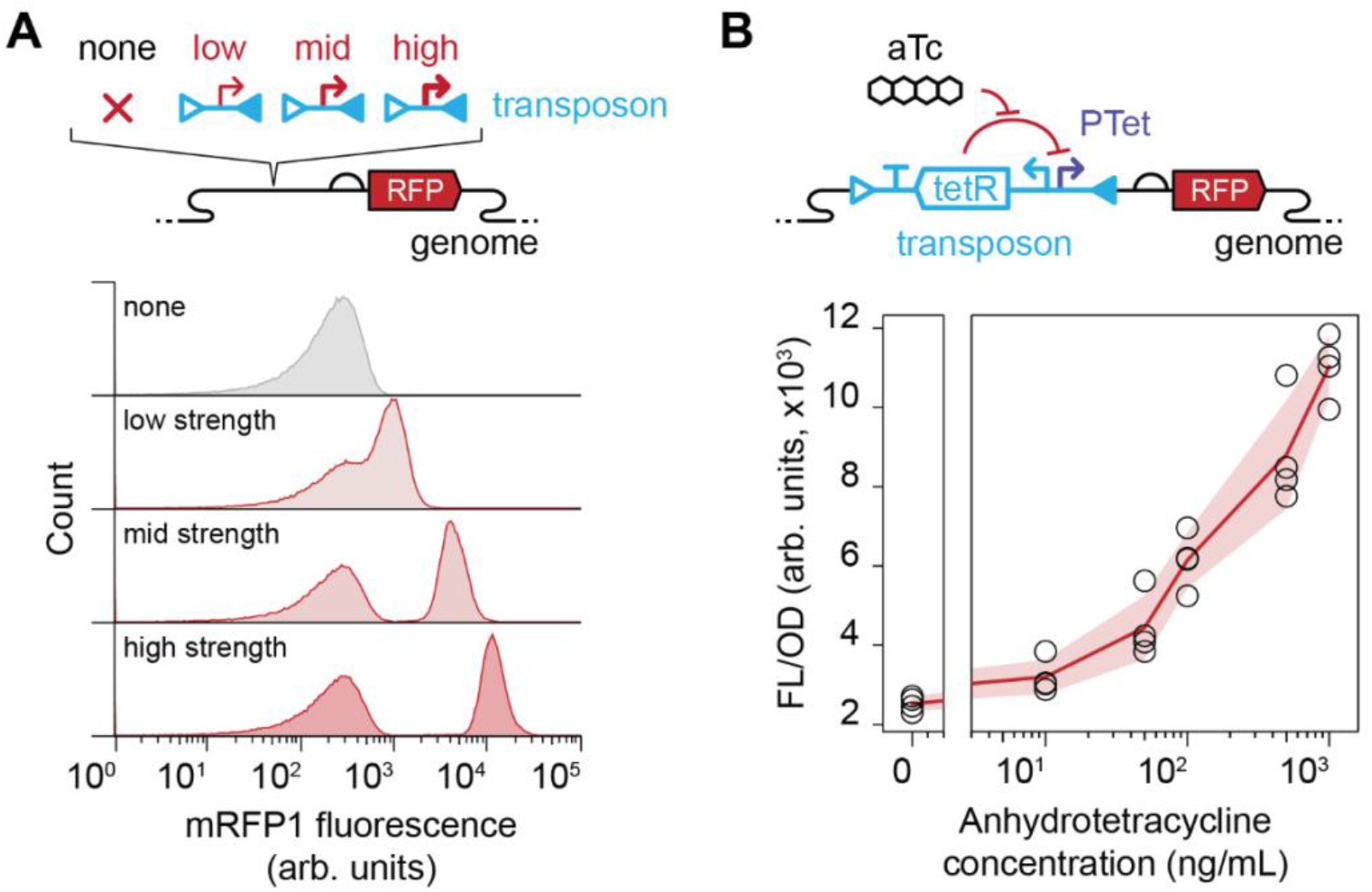
Transcription activation can be tuned by exchanging transposon cargo. (**A**) Schematic of different strength constitutive promoters transposed upstream of mRFP1 using pCAST (top). Flow cytometry data showing single-cell mRFP1 fluorescence of *E. coli* edited with cargos containing high, medium, or low strength constitutive promoters (bottom). Data are representatives of *n* = 3, with the other replicates shown in **Supplementary Fig. 4**. (**B**) Schematic of the PTet inducible promoter system inserted via CAST (top). Bulk fluorescence analysis of *E. coli* cells edited with a transposon containing an inducible promoter cargo (bottom). Data shows fluorescence characterization (measured in units of fluorescence [FL]/optical density [OD] at 600 nm) of mRFP1 observed at varying anhydrotetracycline (aTc) concentrations (bottom). Red line is the average and shading indicates the standard deviation of *n* = 4 biological replicates.

### Activating transcription of endogenous genes via CAST

Having shown transcriptional activation in our reporter strain, our next aim was to test the system within a wild-type (WT) strain for activation of endogenous genes. To do so, we selected four genes dispersed across the *E. coli* MG1655 genome with varying genomic contexts and basal expression levels; *ydiN*, a putative transporter, *zraP*, a signaling pathway modulator, *cadB*, a lysine:cadaverine antiporter, and *ffh*, the protein component of a signal recognition particle (**Figure 3A**).^35^ A library of pCAST plasmids containing a strong constitutive promoter facing the LE direction as a transposon cargo and a crRNA targeting the template strand upstream of each gene was designed and transformed into *E. coli* cells. Then, a PCR screen was conducted to isolate edited strains with transposons in the RE-LE direction. To quantify the expression levels of the targeted genes across these strains, reverse transcription quantitative PCR (RT-qPCR) was performed. This analysis showed that, for all four targeted genes, a significant increase in the expression levels between the WT and the CAST-edited strains was observed, suggesting robust activation of endogenous genes can be achieved with our CAST activation strategy (**Figure 3B**). Interestingly, the activated transcription level achieved in the CAST-edited strains was relatively comparable across the different genes, compared to the greater variance seen in their basal expression levels in the WT strain. This leads to lower fold activations in the case of genes with higher basal expression, such as *ffh*, and suggests that the level of expression in CAST-edited strains is primarily determined by the strength of the promoter cargo inserted. Overall, the results demonstrate the ability to robustly activate the transcription of endogenous genes through our CAST-based promoter insertion strategy.

**Figure 3.**
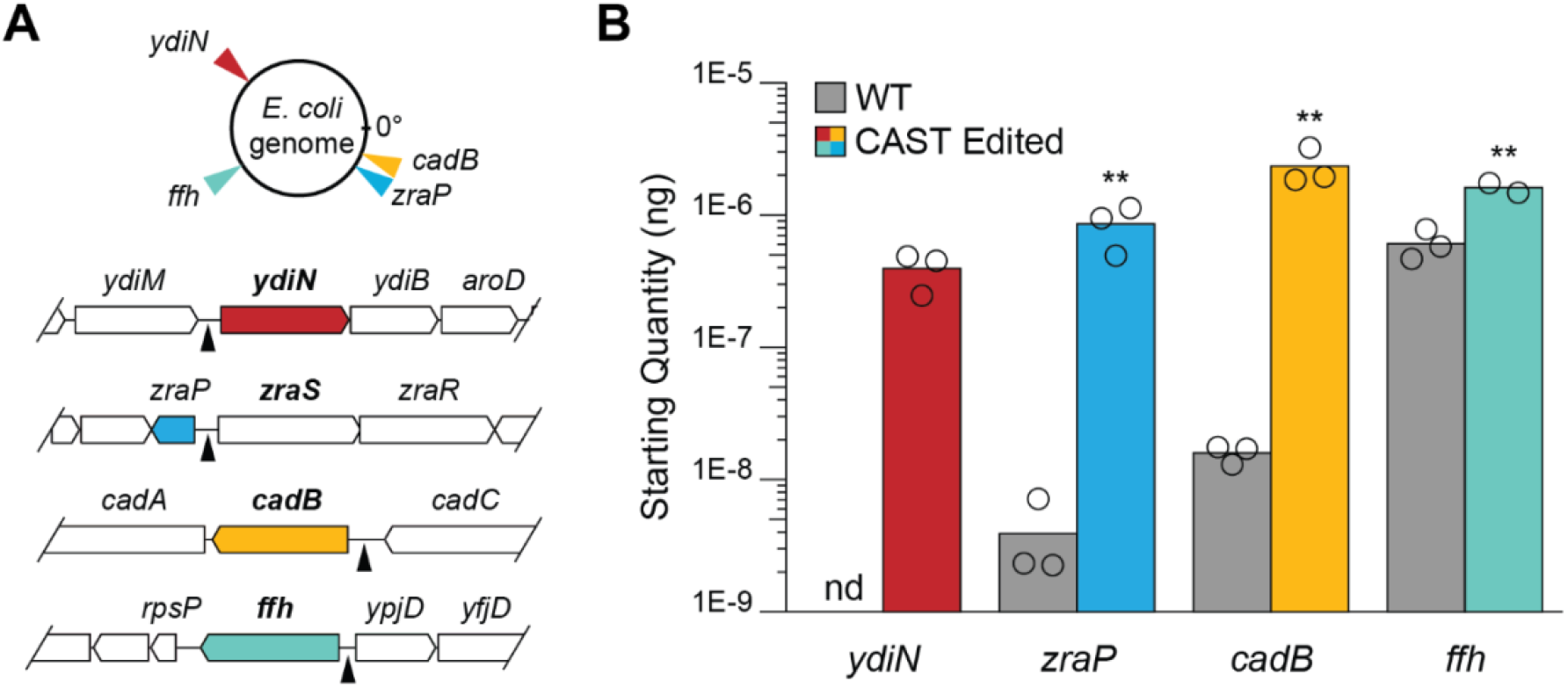
Transcriptional activation of endogenous genes in *E. coli*. (**A**) Location of targeted genes in the *E. coli* K-12 substr. MG1655 genome (top). Genetic context of targeted genes (bottom). Black triangles indicate sites targeted for transposition. (**B**) Starting mRNA quantity of targeted genes with (CAST Edited) and without (WT) insertion of a strong constitutive promoter cargo by the pCAST system, as measured via RT-qPCR. Starting quantities were determined using a relative standard curve. Bars represent averages of *n* = 3 replicates, except for the *ffh* CAST Edited condition that is from *n* = 2 replicates. Statistical significance between conditions is indicated by asterisks (Student’s T-Test, p<0.05). nd = none detected.

### Demonstrating that CAST can selectively activate an ampicillin resistant phenotype

Having demonstrated activation of endogenous genes, we next sought to investigate if we could selectively induce specific phenotypes using CAST activation. To test this, we sought to activate the *ampC* and *marA* genes that have previously been shown to confer increased resistance to the antibiotic ampicillin when overexpressed.^1,3^ As a negative control, we also activated four endogenous genes (*ydiN, zraP, cadB*, and *ffh)* that have no reported effect on ampicillin resistance. pCAST plasmids were designed with the PTet inducible promoter as a transposon cargo, transformed into WT *E. coli*, and incubated at 30°C in liquid culture for 24 hours to allow for transposition. For the *ampC* and *marA* genes, two different crRNA were designed that targeted distinct positions upstream of each gene. Individual colonies were then grown overnight in liquid culture and sub-cultured the next day, followed by PCR analysis to confirm the presence of the intended RE-LE edits in the population (**Supplementary Figure 6**). We note that some of these populations were also detected to contain LE-RE edits and no edits by PCR analysis, which we reasoned would be a small fraction of the population based on our earlier results (**Figure 1C-D**). Finally, the cells were spotted on plates containing or lacking anhydrotetracycline (aTc) to induce expression of the targeted genes as well as varying concentrations of ampicillin to test for growth. From this analysis, we observed that cells where *ampC* and *marA* were targeted by CAST had an enhanced ability to grow at higher concentrations of ampicillin (8 μg/mL) under the induced condition compared to the negative control genes or the uninduced conditions (**Figure 4, Supplementary Figure 7**). This was true for both crRNAs used for *ampC* and *marA*, suggesting that precise targeting upstream of a gene of interest was not necessary for its activation. Overall, the results show that CAST can induce novel phenotypes through activation of endogenous genes.

**Figure 4.**
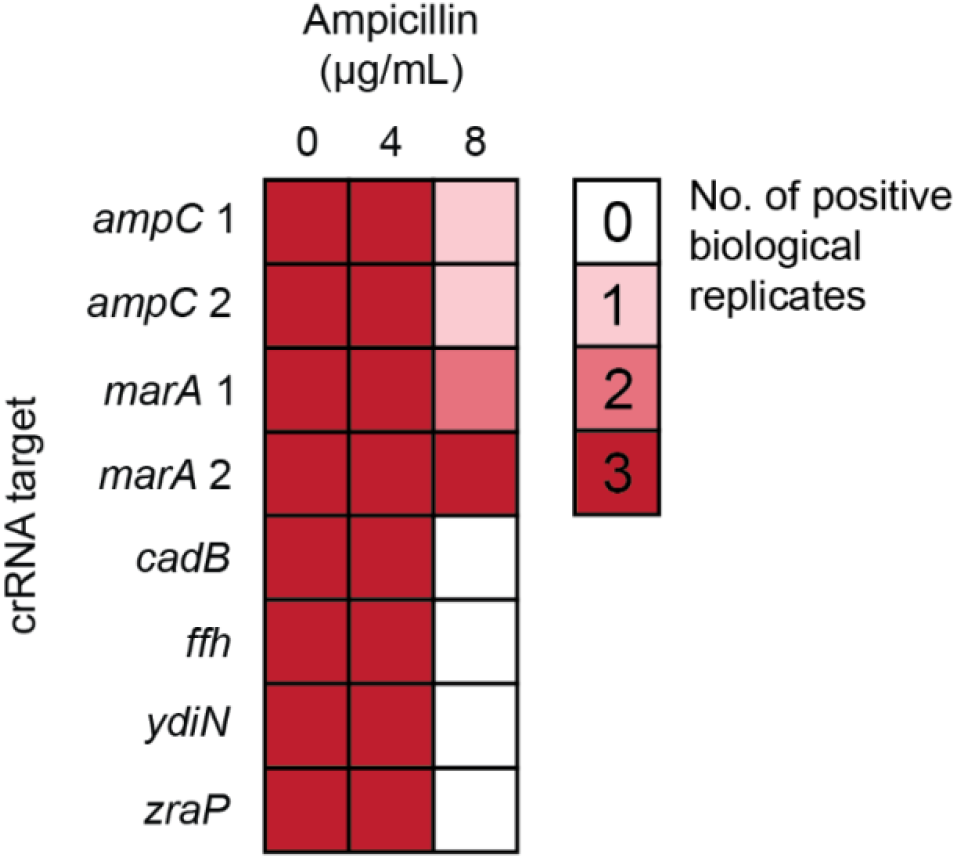
Gain of function via CAST transcriptional activation. The number of positive biological replicates in which growth was observed for different crRNA designs under variable ampicillin concentrations. Full data is shown in **Supplementary Fig. 7**. For each biological replicate, three technical replicates were performed. For a biological replicate to be considered positive, growth had to be observed in >2 technical replicates; otherwise, they were scored as having no growth.

## Discussion

In this work, we demonstrated the use of the *V. cholerae* CAST system for transcriptional activation. We first showed activation of an mRFP1 target gene within a population and then tuned the level of expression using variable strength and inducible promoter elements. We reported high level of editing and activation efficiency, which we believe could be further increased through further optimizing of experimental conditions.^33^ Our work also showed robust activation of endogenous *E. coli* genes through the insertion of strong constitutive promoters; Because the inserted promoter was stronger than the native promoters, the genes chosen all showed a significant increase in expression. Finally, we activated the *ampC* and *marA* genes in a WT *E. coli* to induce an ampicillin-resistant phenotype, showcasing the potential for CAST activation to be used as a method for studying genotype-phenotype interactions. Overall, we present a novel CRISPR-based strategy for targeted gene activation.

Within the field of microbiology and synthetic biology, the perturbation of endogenous gene expression is a broadly useful capability. We anticipate that the CAST system will be particularly well-suited for genome-wide expression perturbation studies, as now both knock-outs,^29–31^ and knock-ups of gene expression through the CAST system are shown to be possible. Genome-wide, bidirectional studies have been highly successful in eukaryotes for uncovering gene function through both the repression and overexpression of individual genes.^11^ The CAST system is well-suited to perform similar screens in bacteria due to several features. For one, the ability to use the same transposon for different insertion sites means only the crRNA needs to be varied, making it more suitable for library generation through pooled DNA synthesis. Furthermore, the *V. cholerae* CAST has a highly promiscuous protospacer-adjacent motif (PAM) preference, with a general bias for a 5’–CN–3’ PAM but high flexibility to other sequences, expanding the possible targetable sites across a genome.^36^ These characteristics distinguish the CAST system from other CRISPR-based technologies, making it an attractive activation strategy for high-throughput gene editing.

While this work has focused on studies in *E. coli*, we anticipate it is possible to expand this method to other microbes. For example, the VchCAST system has been adapted for use in both gram-negative and -positive species, including biotechnologically relevant species such as *Lactococcus lactis*^37^, *Corynebacterium glutamicum*, and *Bacillus subtilis*.^38^ This CAST system has also been used for genome editing of non-model species within soil and gut microbial communities.^31^ Additionally, bioinformatic analysis of genomes has uncovered a diversity of CAST systems that could be adapted for our approach. This includes the commonly used *Scytonema hofmanni* type V-K CAST,^39^ a large variety of type I-F CAST systems,^34,40,41^ as well as some type I-B and I-D CAST systems.^42–44^ These and other systems can provide alternatives with different PAM preferences, transposition mechanisms, and insertion behaviors, expanding the possibilities for applying CAST systems as transcriptional perturbators.

In summary, we present a novel approach for gene activation that leverages the unique RNA-guided transposition of CAST systems. We anticipate that our newly described tool is likely to assist efforts to both fundamentally understand and engineer microbes in the coming years.

## Materials and Methods

### Plasmid assembly and strain engineering

All plasmids and strains used in this study can be found in **Supplementary Tables 1** and **2**, respectively, and example annotated plasmids and strains can be found in **Supplementary Table 3**. Plasmids containing the CAST system were derived from pSL1142 (Addgene plasmid #160730).^33^ All experiments were performed in wild-type (WT) *E. coli* str. K-12 substr. MG1655 or engineered derivatives. The sfGFP-mRFP1 reporter strain (sJEC042), and the versions of it with a promoter insert facing sfGFP (sJEC050) or mRFP1 (sJEC043), were created using CRISPR-Cas9 genome editing.^22^

### Genome editing using CAST transposition

All crRNAs used in this study can be found in **Supplementary Table 4**. Plasmids were transformed into chemically competent *E. coli* cells, plated on LB-Agar with appropriate antibiotics (LB-agar-Ab), and incubated at 37°C for 24 hours. The plates were then resuspended in 3 mL LB media with antibiotics (LB-Ab) and 100 μL of the suspension was used to inoculate 10 mL of fresh LB-Ab. These cultures were then incubated at 30°C shaking at 200-255 rpm for 24 hours. For experiments using variable strength promoters, the plate suspension was used to inoculate 10 mL of fresh LB-Ab to an optical density at 600 nm (OD) of ∼0.05 and incubated at 37°C shaking at 200-255 rpm for 24 hours. A similar protocol was followed for creating the sfGFP-mRFP1 reporter strains with the PTet inducible promoter transposon (sJEC057), and the WT strains with the PTet inducible promoter transposon for ampicillin experiments (pJEC1258-65) or the strong constitutive promoter transposon for RT-qPCR analysis (sJEC051-4). The post-incubation populations were plated on LB-agar-Ab and incubated overnight at 37°C to obtain single colonies. For experiments using sJEC051-4 and sJEC057, a clonal strain containing the desired insert in the genome was identified by PCR and confirmed by sequencing, and then used for fluorescence experiments or RT-qPCR analysis. For experiments using pJEC1258-65, colonies were randomly selected and used for the ampicillin resistance experiments.

### Flow cytometry measurement and analysis

Flow cytometry was performed on the Sony Biotechnology SH800 Cell Sorter. A total of 100,000 events were measured for each biological replicate. FCS files were analyzed using the FlowJo software. Each condition was gated to contain >90% of the measured events, and this sub-population was used for analysis. Fluorescence compensation was performed using samples containing the sfGFP-mRFP1 reporter strain (sJEC042) with a non-targeting plasmid as a negative control and the sfGFP-mRFP1 reporter strain edited to contain the cargo with the constitutive promoter driving either mRFP1 (sJEC043) or sfGFP (sJEC050) expression as positive controls. The quadrant gates used for Fig. 1C-D were manually set at 1,000 arb. units on FlowJo.

### Characterization of insertion distance by amplicon sequencing

Fluorescence Assisted Cell Sorting was performed on a Sony Biotechnology SH800 Cell Sorter. A total of 500,000 events per biological replicate were sorted using the gates shown in **Supplementary Figure 2** into 5 mL LB. The cultures were incubated at 37°C shaking at 200-255 rpm for 1 hour, followed by the addition of 15 mL LB-Ab and further incubation at 37°C shaking at 200-255 rpm overnight. The Wizard Genomic DNA Purification Kit (Promega) was used for genome extraction. PCR amplification of the region between the left transposon end and the 5’ end of mRFP1 was performed for each of the replicates, column purified, and amplicon sequencing performed (Amplicon-EZ, Genewiz). Primers used can be found in **Supplementary Table 5**. From this data, the distance between the 3’ end of the crRNA binding site and the 5’ end of the transposon was determined, accounting for the 5 bp duplication that results after insertion.

### Bulk fluorescence measurement and analysis

Chemically competent *E. coli* were transformed with plasmids or streaked out from glycerol stocks onto LB-Agar-Ab, and then incubated overnight at 37°C. For each replicate, individual colonies were used to inoculate 300 μL of LB-Ab in a deep 96-well plate and incubated overnight at 37°C shaking at 1,000 rpm. Subsequently, 5 μL of overnight culture was used to inoculate 295 μL of fresh, pre-warmed LB-Ab which was then incubated at 37°C shaking at 1,000 rpm for 4 hours. For experiments using inducible promoter donors, 5 μL of overnight culture was used to inoculate 290 μL of fresh, pre-warmed LB-Ab which was then incubated at 37°C shaking at 1,000 rpm for 4 hours. The cultures were then induced by adding 5 μL of anhydrotetracycline (Cayman Chem) to final concentrations of 10, 50, 100, 500, and 1,000 ng/mL, and incubated a further 4 hours at 37°C shaking at 1,000 rpm.

For bulk fluorescence measurements, a 96 well plate with 25 μL of culture in 75 μL of phosphate buffer saline (PBS) was used. A media-only control of 25 μL LB and 75 μL PBS was also included. Optical density at 600 nm (OD) was measured, as was fluorescence (FL) for sfGFP (485 excitation, 535 emission) and mRFP1 (560 nm excitation, 590 nm emission). For analysis, the mean OD and FL of the media control was subtracted from each well. The ratio of FL to OD was then measured for each well.

### Total RNA extraction

*E. coli* cells were streaked out from glycerol stocks on LB-agar-Ab and incubated overnight at 37°C. Colonies were used to inoculate LB-Ab and incubated overnight at 37°C shaking at 200-255 rpm. 10 mL of fresh, pre-warmed LB-Ab was inoculated with 100 μL of overnight culture and grown at 37°C shaking at 200-255 rpm to an OD of 0.4-0.6. Total RNA was extracted using the Quick RNA Miniprep Plus Kit (Zymo Research) following the manufacturer’s instructions for isolating large RNA species and without in-column DNase treatment. Total RNA was then measured on a Qubit fluorometer (Invitrogen). DNase treatment was performed using Turbo DNase (Invitrogen), treating ∼300 ng of total RNA with 4 U of enzyme, and incubating at 37°C for 1 hour. Reaction clean-up was then performed using the Quick RNA Miniprep Plus Kit, again following the manufacturer’s instructions and without DNase treatment.

### Reverse transcription quantitative PCR (RT-qPCR)

Total RNA samples were measured on a Qubit fluorometer and normalized to a concentration of 0.5 ng/μL. Reverse transcription was performed by mixing 3.5 μL of water, 0.5 μL of 2 μM RT primer, 0.5 μL of 10 mM dNTPs, and 2 μL of 0.5 ng/μL total RNA. This mixture was incubated at 65°C for 5 min followed by incubation on ice for a further 5 mins. A mixture of 0.5 μL 100 mM Dithiothreitol (DDT), 2 μL 5X First-Strand Buffer (Invitrogen), 0.5 μL 40 U/μL RNase OUT (Invitrogen), 0.5 μL 200 U/μL SuperScript III Reverse Transcriptase (Invitrogen), and the pre-mixture were combined. The reaction was then incubated at 55°C for 1 hour, followed by 70°C for 15 min, and finally stored at -80°C. A CFX Connect Real-Time PCR system (Bio-Rad) was used for qPCR analysis. Reactions were performed using a 96 well plate with an optically clear microseal ‘B’ film (Bio-Rad). Three technical replicate reactions were performed for each biological replicate. Each reaction contained 0.5 μL of each 2 μM qPCR primer stock, 5 μL of 2X Maxima SYBR Green/ROX qPCR Master Mix (Thermo Scientific), 3 μL of water, and 1 μL of the RT product. To confirm digestion of DNA during the DNase treatment, qPCR analysis was performed directly on the total RNA (i.e., a no-RT control). Additionally, no-template control reaction was also performed using 1 μL of water in place of the RT product. For each qPCR primer set, a standard curve was created using serial dilution of PCR-amplified products. Reaction protocol was as follows: 50°C for 2 min, 95°C for 10 min, and 35 cycles of 95°C for 15 sec, 60°C for 11 min, and measurement of fluorescence. Melting curve analysis was performed after amplification to confirm the absence of primer dimers. RT and qPCR primers used are listed in **Supplementary Table 4**.

For data analysis, CFX Manager Software (Bio-Rad) was used to determine Cq values. To determine the starting quantities (SQ) of each gene of interest, a standard curve was used. In brief, the mean Cq of the technical replicates was determined for each dilution of the gene standard. This mean was fitted to *y = mx + b*, with *y* being the mean Cq values and *x* the log SQ. The formula was then used to calculate the SQ of each technical replicate for the test samples, and the technical replicate SQ values were then averaged to obtain a biological replicate SQ.

### Ampicillin resistance experiments

For each biological replicate, individual colonies were used to inoculate 300 μL of LB-Ab in a deep 96-well plate and incubated overnight at 37°C shaking at 1,000 rpm. The next day, 5 μL of overnight culture was used to inoculate 295 μL of fresh, pre-warmed LB-Ab and incubated at 37°C shaking at 1,000 rpm for 4 hours. The OD of each sub-culture was measured as outlined in “bulk fluorescence measurement and analysis,” and found to be ∼0.6-0.8. A one-way ANOVA was performed on the OD values to confirm equal growth across the conditions. Samples were also taken from each sub-culture for the PCR analysis in **Supplementary Figure 6**. Each biological replicate was then spotted 3 times, for technical replicates, on LB-agar-Ab plates with 0 μg/mL, 4 μg/mL, or 8 μg/mL ampicillin (A9518, Sigma-Aldrich) and with or without 1000 ng/mL anhydrotetracycline, and then incubated for 48 hr at 37°C. For each biological replicate, growth on at least two out of the three technical replicates was counted as a positive hit.

## Supporting Information

Additional experimental details and measurements; and plasmids, strains, and primers used in this study (DOCX)

## Author Contribution

A.G. and J.C designed the study and experiments, and prepared the manuscript. A.G. collected all experimental data.

## Conflict of Interest

The authors declare no competing interests.

## Acknowledgements

The authors acknowledge the Chappell Lab members for helpful discussion. The authors also wish to acknowledge Daniel J. Haller for his assistance in the design and cloning of sJEC057, as well as Biki B. Kundu for his expertise in CRISPR-Cas9 editing that aided the creation of the reporter strains used in this study. This material is based on work supported by the National Science Foundation (grant no. # 2237512).

## Notes

### Competing Interest Statement

The authors have declared no competing interest.

