## Supplemental Information for "Targeted Transcriptional Activation using a CRISPR-Associated Transposon System"

### Table of Contents


| Title | Page(s) |
|---|---|
| Supplementary Figure 1. Thresholds used to determine mRFP1 and sfGFP positive populations in flow cytometry analysis | 2 |
| Supplementary figure 2. Determining the distance between the crRNA target site and transposon insertion | 3 |
| Supplementary Figure 3. Bulk fluorescence analysis of the effect of pCAST expression on transcription of transposon activated genes | 4 |
| Supplementary Figure 4. Single-cell fluorescence analysis of *E. coli* cells edited with pCAST plasmids containing variable strength promoters as transposon cargo | 5 |
| Supplementary Figure 5. Bulk Fluorescence analysis of *E. coli* cells edited with a pCAST plasmid containing an inducible promoter as transposon cargo | 6 |
| Supplementary Figure 6. PCR confirmation of CAST edits | 7 |
| Supplementary Figure 7. Growth patterns of *E. coli* populations edited with pCAST to confer antibiotic resistance | 8 |
| Supplementary Table 1. Plasmids used in this study | 9 |
| Supplementary Table 2. Strains used in this study | 10 |
| Supplementary Table 3. Sample plasmids and strains | 11-30 |
| Supplementary Table 4. crRNAs used in this study | 31 |
| Supplementary Table 5. Primers used in this study | 32 |

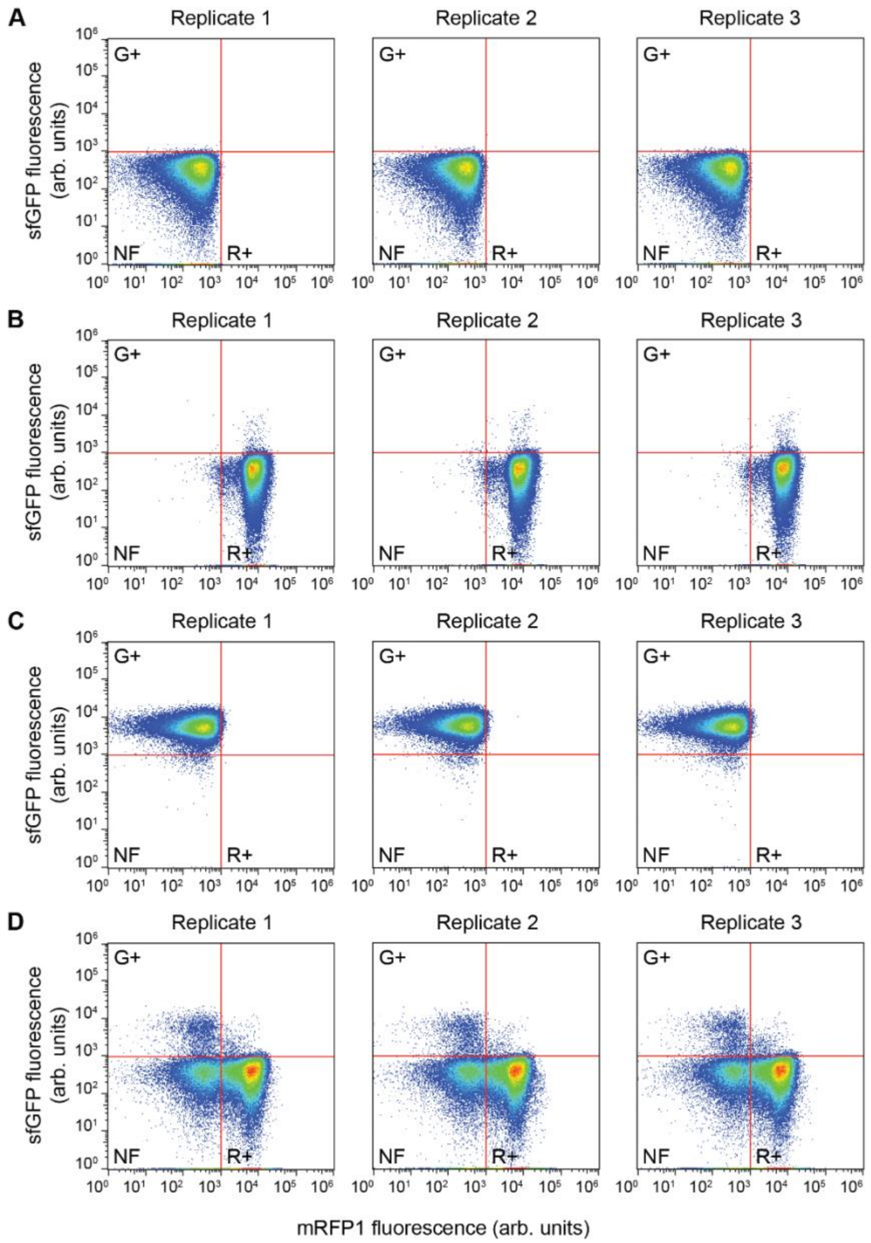


**Supplementary Figure 1. Thresholds used to determine mRFP1 and sfGFP positive populations in flow cytometry analysis.** Flow cytometry data showing the single-cell sfGFP and mRFP1 fluorescence of different populations of *E. coli* cells. (**A**) Data from the sfGFP-mRFP1 reporter strain transformed with a non-targeting pCAST plasmid. (**B**) Data from the sfGFP-mRFP1 reporter strain edited to contain a transposon insert with the constitutive promoter driving mRFP1 expression (RE-LE), transformed with a targeting pCAST plasmid. (**C**) Data from the sfGFP-mRFP1 reporter strain edited to contain a transposon insert with the constitutive promoter driving sfGFP expression (LE-RE), transformed with a targeting pCAST plasmid. (**D**) Data from the sfGFP-mRFP1 reporter strain transformed with a targeting pCAST plasmid. Three biological replicates are included for each condition with replicates 1 being presented in Figure 1C. The thresholds used to assign sfGFP and mRFP1 positive populations are indicated on each panel.

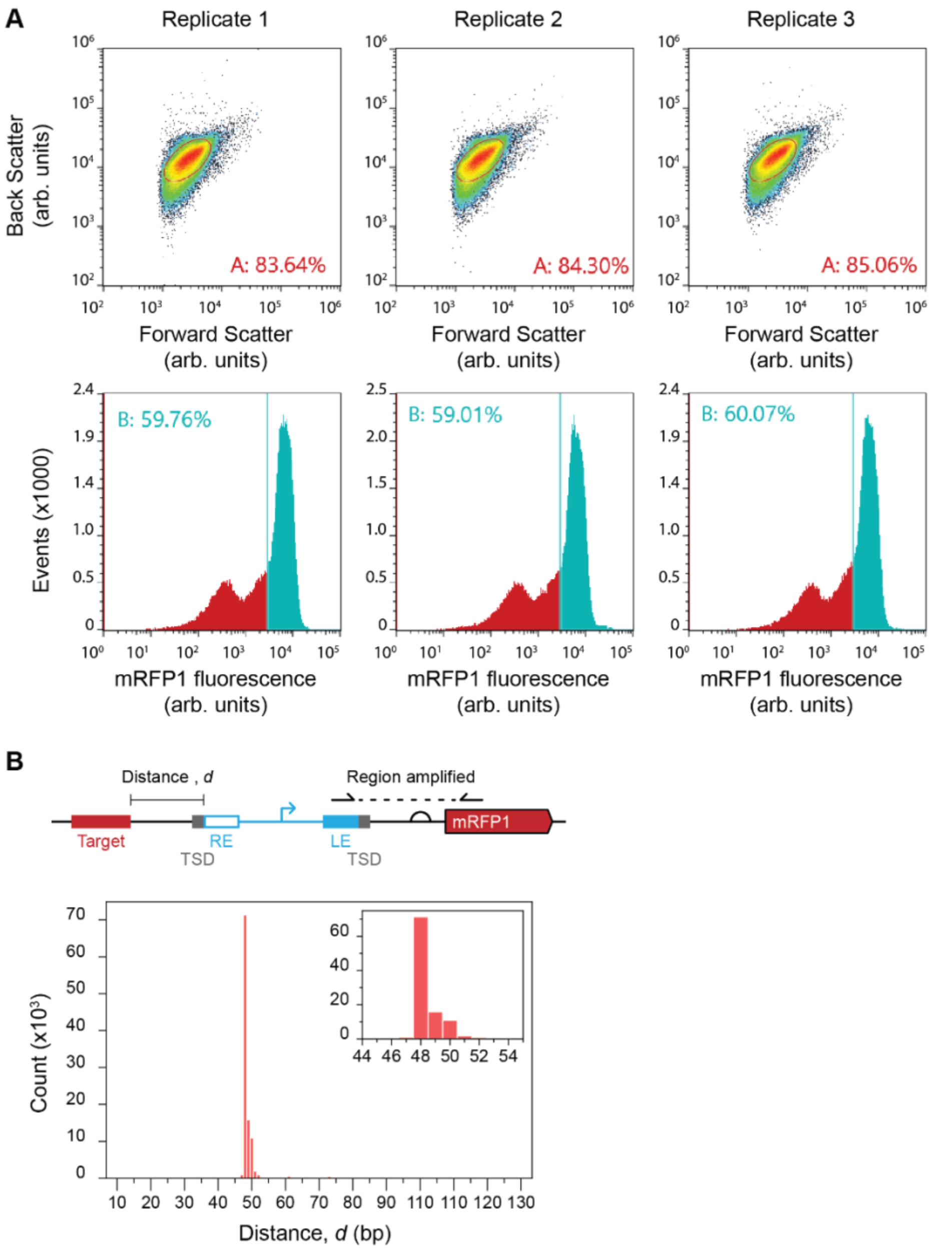


**Supplementary figure 2. Determining the distance between the crRNA target site and transposon insertion.** (**A**) Gating used for fluorescence activated cell sorting (FACS) of mRFP1 positive populations. (Top panel) The sfGFP-mRFP1 reporter strain transformed with a targeting pCAST plasmid. Populations from Fig. 1C-D were first gated using the forward and back scatter to capture the majority of the cell population (gate "A"). (Bottom panel) A second gate was then applied to capture mRFP1 positive cells (bottom, gate "B"). (**B**) (Top panel) Schematic of the distance (*d*) between the 3' end of the crRNA binding site and the 5' end of the transposon insertion position along with the PCR primer sites used for amplicon sequencing. Distance (*d*) includes the 5 bp target site duplication (TSD) generated during insertion. (Bottom panel) Quantification of *d* using amplicon sequencing of PCR product, with insert showing counts within the 44-55 bp range.

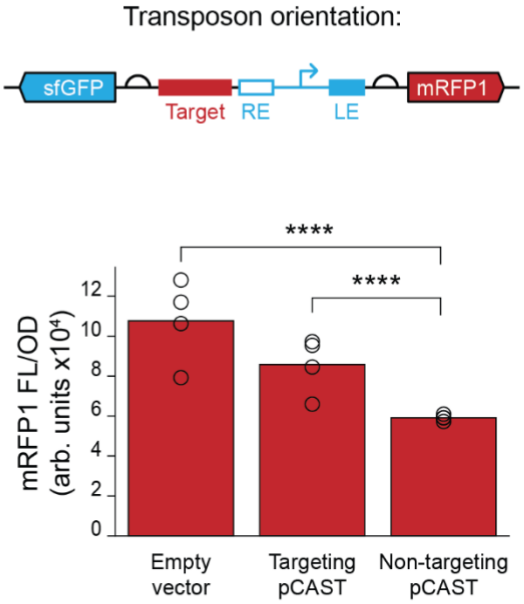


**Supplementary Figure 3. Bulk fluorescence analysis of the effect of pCAST expression on transcription of transposon activated genes.** The sfGFP-mRFP1 reporter strain edited to contain a transposon insert with the constitutive promoter driving mRFP1 expression (RE-LE) were transformed with a non-targeting pCAST plasmid, a targeting pCAST plasmid, or an empty vector control. Data shows fluorescence characterization (measured in units of fluorescence [FL]/optical density [OD] at 600 nm) for mRFP1 of the varying conditions. Bars are averages of $n$ = 3. Statistical significance between populations is indicated by asterisks (Student's T-Test **** = $p<0.0005$).

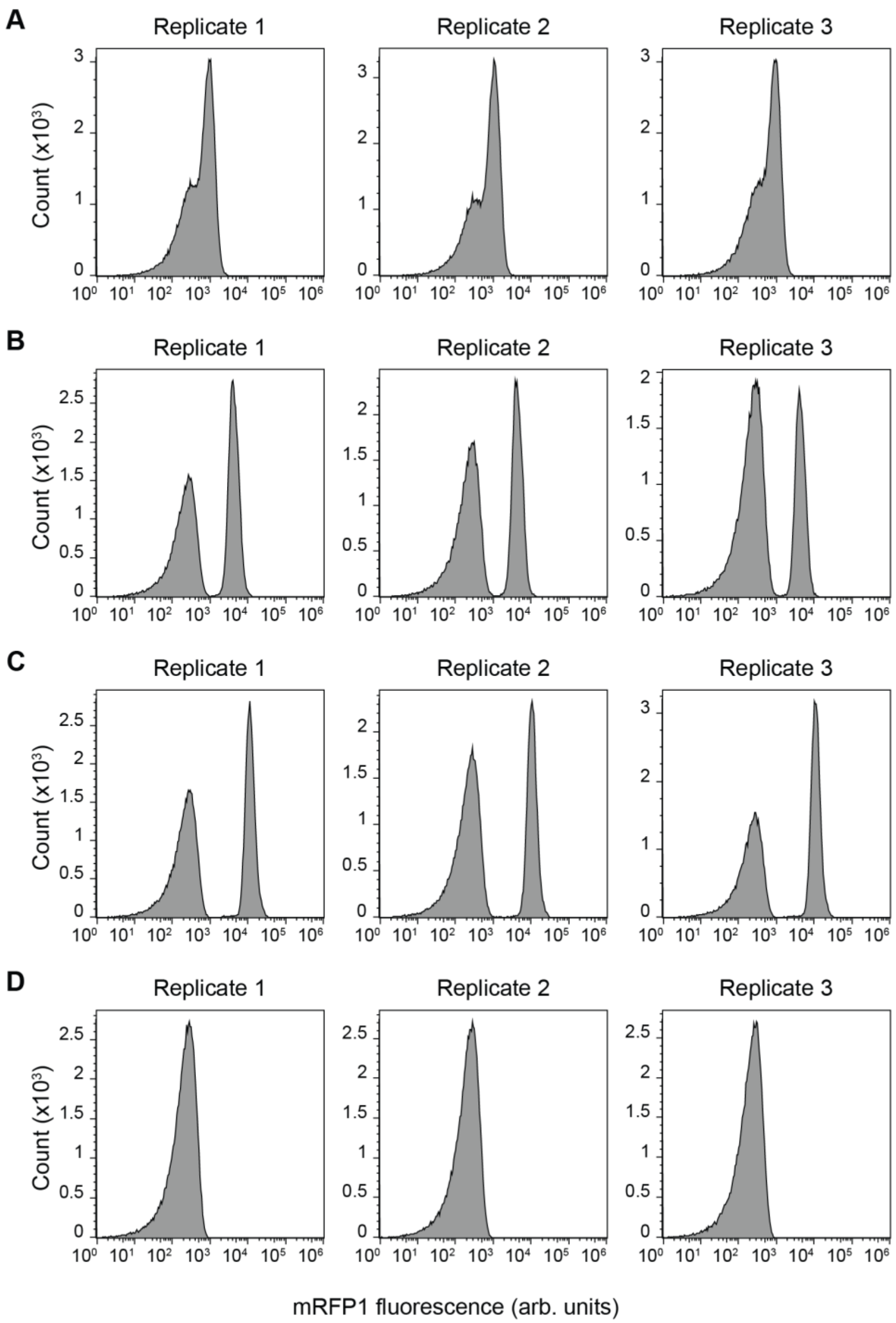


**Supplementary Figure 4. Single-cell fluorescence analysis of *E. coli* cells edited with pCAST plasmids containing variable strength promoters as transposon cargo.** Flow cytometry data showing single-cell mRFP1 fluorescence of different populations of *E. coli* cells. Data from the sfGFP-mRFP1 reporter strain edited with a pCAST plasmid containing a (A) low strength, (B) medium strength, or (C) high strength promoter as transposon cargo. (D) Data from the sfGFP-mRFP1 reporter strain transformed with a non-targeting pCAST plasmid. Three biological replicates are included for each condition with replicates 1 being presented in Figure 1C.

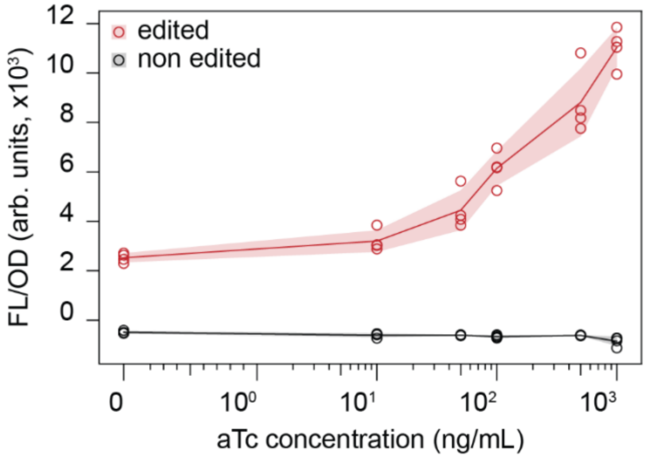


**Supplementary Figure 5. Bulk Fluorescence analysis of *E. coli* cells edited with a pCAST plasmid containing an inducible promoter as transposon cargo.** Data shows fluorescence characterization (measured in units of fluorescence [FL]/optical density [OD] at 600 nm) of mRFP1 observed at varying anhydrotetracycline (aTc) concentrations in the sfGFP-mRFP reporter strain with (red points) or without (black points) editing with the inducible promoter donor. Lines are the average and shading indicates standard deviation of $n$ = 4 biological replicates.

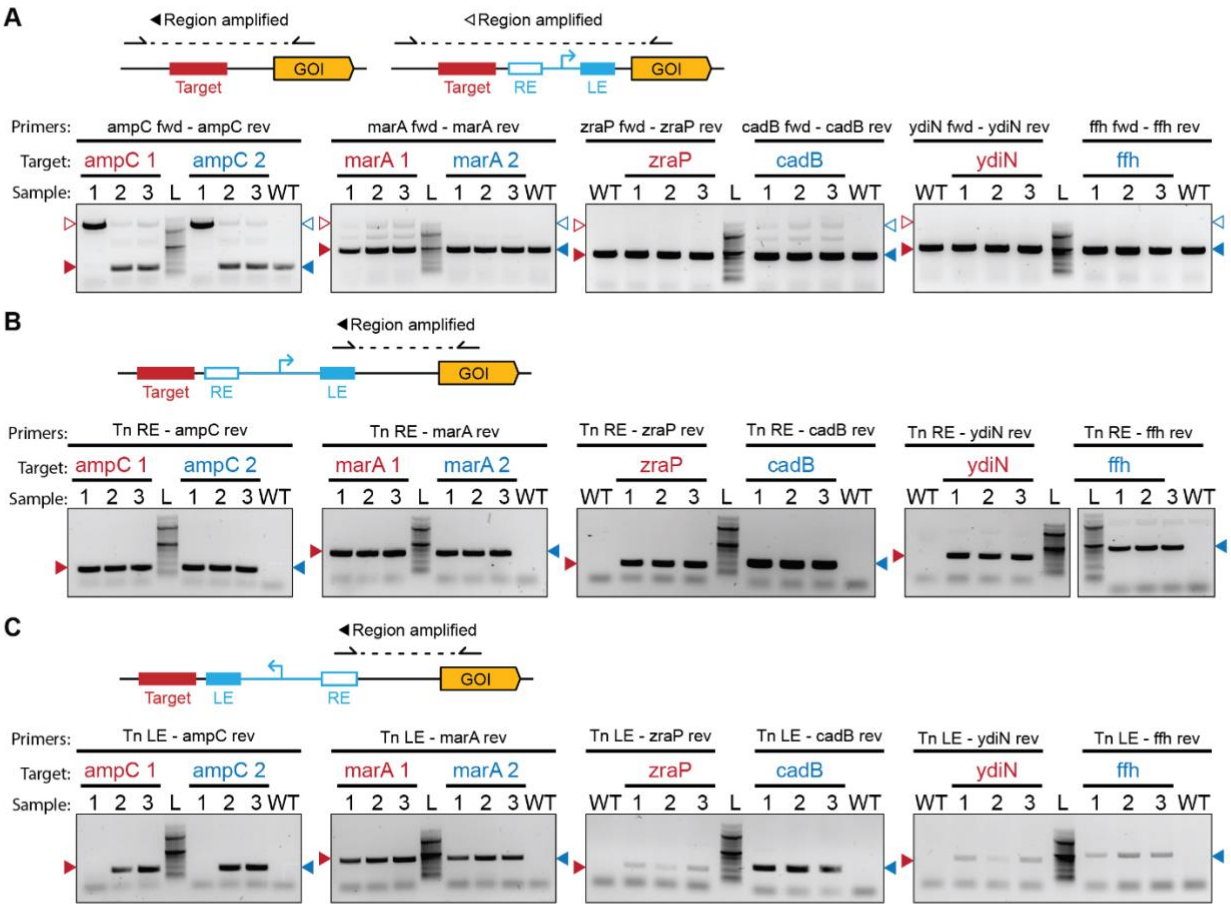


**Supplementary Figure 6. PCR confirmation of CAST edits.** Gel electrophoresis of PCR products from wild-type *E. coli* populations incubated with a pCAST plasmid containing a crRNA for the *ampC*, *marA*, *cadB*, *ffh*, *ydiN*, or *zraP* genes, and an inducible PTet promoter system as transposon cargo. (**A**) Schematic of the genomic locus with and without transposon insertions amplified by the PCR (top) and gel electrophoresis of PCR products from the amplification (bottom). Shaded triangles indicate products resulting from a lack of transposon insertions while open triangles indicate products resulting from either RE-LE or LE-RE oriented insertions. Triangle colors indicate the target. Samples are labelled numerically for each biological replicate, "L" for the DNA ladder (NEB 100 bp Ladder, N3231), and "WT" for PCR products where a colony of *E. coli* transformed with a non-targeting version of the pCAST plasmid containing the same cargo was used as a template. Schematic of the genomic locus with RE-LE oriented (**B**) or LE-RE oriented (**C**) transposon insertions amplified by the PCR (top) and gel electrophoresis of PCR products from the amplification (bottom), as in A. We note that due to amplification bias for shorter DNA products, in panel A, for some samples we did not observe the amplification of transposon insertions that were expected based on the results shown in B and C.

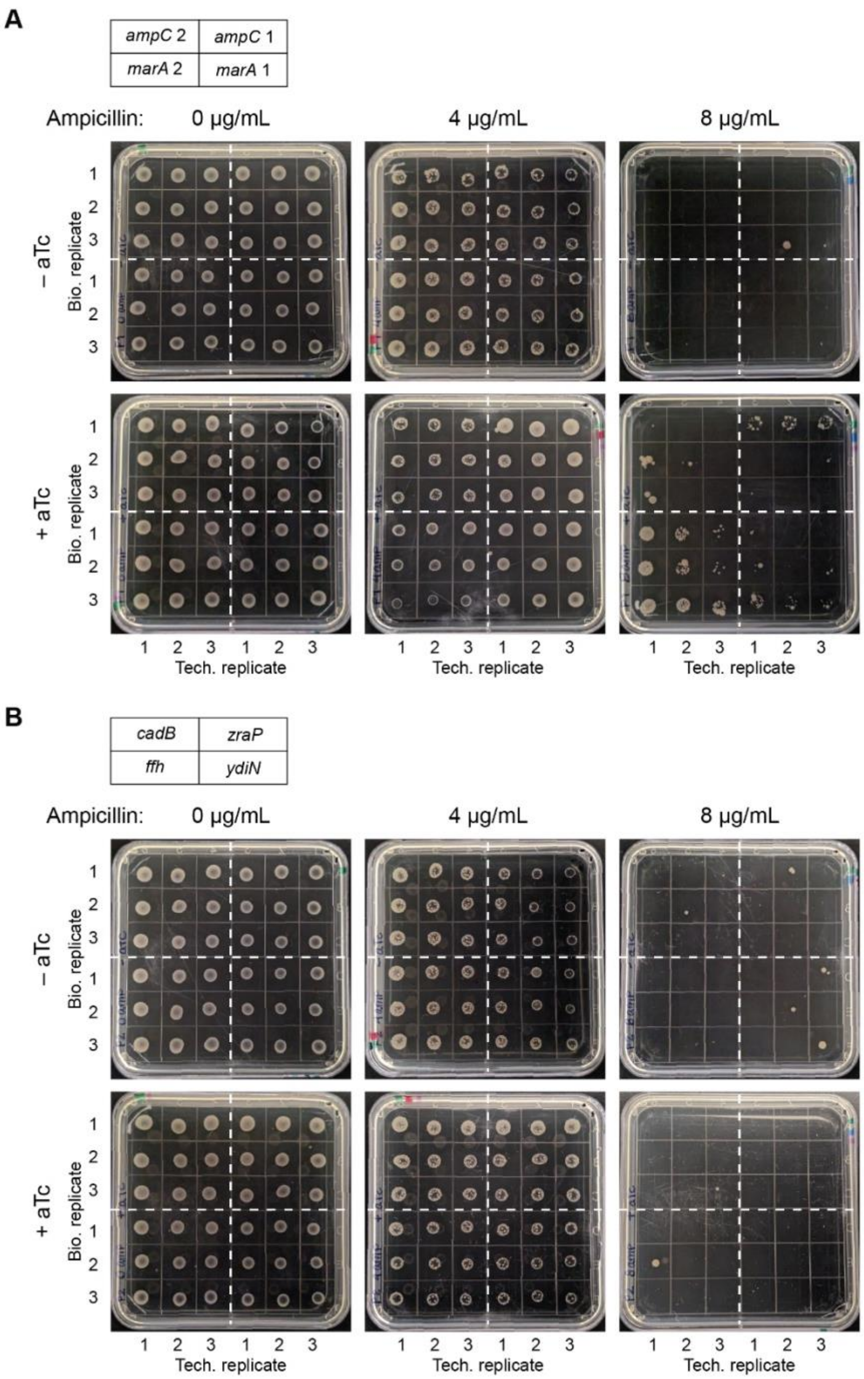


**Supplementary Figure 7. Growth patterns of *E. coli* populations edited with pCAST to confer antibiotic resistance.** (**A**) Legend for plates indicating the crRNA used for targeting insertion of the PTet inducible promoter cargo into the *ampC* and *marA* genes (top). Plates with or without anhydrotetracycline (aTc) inducer and with varying ampicillin concentrations after 48 hr incubation at 37°C (bottom). (**B**) Legend for plates indicating the crRNA used for targeting insertion of the PTet inducible promoter cargo into the *cadB*, *ffh*, *zraP*, and *ydiN* genes (top). Plates with or without anhydrotetracycline (aTc) inducer and with varying ampicillin concentrations after 48 hr incubation at 37°C (bottom).

| Supplementary Table 1. Plasmids used in this study. | | | |
|---|---|---|---|
| **Name** | **Description** | **crRNA** | **Used for figures** |
| pJEC121 | pCOLADuet empty vector | | SF3 |
| pJEC1249 | pCAST, non-targeting crRNA, J23119 cargo | non-targeting | 1C, 1D, 2A, SF1, SF3, SF4, SF5 |
| pJEC1250 | pCAST, mRFP1 crRNA, J23119 cargo | mRFP1 | 1C, 1D, 2A, SF1, SF2, SF3, SF4 |
| pJEC1251 | pCAST, mRFP1 crRNA, J23101 cargo | mRFP1 | 2A, SF4 |
| pJEC1252 | pCAST, mRFP1 crRNA, J23105 cargo | mRFP1 | 2A, SF4 |
| pJEC1253 | pCAST, zraP crRNA, J23119 cargo | zraP | (to make sJEC051 for 3B) |
| pJEC1254 | pCAST, cadB crRNA, J23119 cargo | cadB | (to make sJEC052 for 3B) |
| pJEC1255 | pCAST, ydiN crRNA, J23119 cargo | ydiN | (to make sJEC053 for 3B) |
| pJEC1256 | pCAST, ffh crRNA, J23119 cargo | ffh | (to make sJEC054 for 3B) |
| pJEC1257 | pCAST, mRFP1 crRNA, PTet cargo | mRFP1 | (to make sJEC057 for 2B) |
| pJEC1258 | pCAST, ampC 1 crRNA, PTet cargo | ampC 1 | 4, SF6, SF7 |
| pJEC1259 | pCAST, ampC 2 crRNA, PTet cargo | ampC 2 | 4, SF6, SF7 |
| pJEC1260 | pCAST, marA 1 crRNA, PTet cargo | marA 1 | 4, SF6, SF7 |
| pJEC1261 | pCAST, marA 2 crRNA, PTet cargo | marA 2 | 4, SF6, SF7 |
| pJEC1262 | pCAST, cadB crRNA, PTet cargo | cadB | 4, SF6, SF7 |
| pJEC1263 | pCAST, ffh crRNA, PTet cargo | ffh | 4, SF6, SF7 |
| pJEC1264 | pCAST, ydiN crRNA, PTet cargo | ydiN | 4, SF6, SF7 |
| pJEC1265 | pCAST, zraP crRNA, PTet cargo | zraP | 4, SF6, SF7 |
| pJEC1266 | pCAST, no crRNA, PTet cargo | | SF6 |

| Supplementary Table 2. Strains used in this study. | | | |
|---|---|---|---|
| **ID** | **Description** | **Plasmids in strain** | **Used for figures** |
| sJEC042 | sfGFP-mRFP1 reporter strain | none | 1C, 1D, 2A, SF1, SF2, SF4, SF5 |
| sJEC081 | sJEC042, RE-LE transposon (J23119 cargo) cloned upstream of mRFP1 | none | SF1, SF3 |
| sJEC050 | sJEC042, LE-RE transposon (J23119 cargo) cloned upstream of mRFP1 | none | SF1 |
| sJEC051 | MG1655, RE-LE transposon (J23119 cargo) CAST-inserted upstream of zraP | JEC1253 | 3B |
| sJEC052 | MG1655, RE-LE transposon (J23119 cargo) CAST-inserted upstream of cadB | JEC1254 | 3B |
| sJEC053 | MG1655, RE-LE transposon (J23119 cargo) CAST-inserted upstream of ydiN | JEC1255 | 3B |
| sJEC054 | MG1655, RE-LE transposon (J23119 cargo) CAST-inserted upstream of ffh | JEC1256 | 3B |
| sJEC055 | MG1655 with a non-targeting pCAST | JEC1249 | 3B |
| sJEC057 | sJEC042, RE-LE transposon (PTet cargo) CAST-inserted upstream of mRFP1 | JEC1257 | 2B, SF5 |

**Supplementary Table 3. Sample plasmids and strains.**

**pJEC1249**

```
LOCUS       pJEC1249        13898 bp ds-DNA     circular
FEATURES             Location/Qualifiers
     promoter        1..35
                     /label="J23119"
     misc_feature    59..86
                     /label="CRISPR repeat"
     misc_feature    complement(87..92)
                     /label="BsaI"
     misc_feature    93..112
                     /label="Ext BsaI"
     misc_feature    113..118
                     /label="BsaI"
     misc_feature    119..146
                     /label="CRISPR repeat"
     RBS             164..169
                     /label="RBS"
     CDS             178..1362
                     /label="TniQ"
     CDS             1363..3285
                     /label="Cas8"
     CDS             3257..4315
                     /label="cas7/csy3"
     CDS             4318..4917
                     /label="cas6f"
     RBS             4926..4931
                     /label="RBS"
     CDS             4939..5631
                     /label="tnsA"
     CDS             5624..7435
                     /label="tnsB"
     CDS             7447..8439
                     /label="tnsC"
     terminator      8477..8524
                     /label="T7 terminator"
     misc_feature    8545..8552
                     /label="Transposon site duplication"
     misc_feature    8545..8655
                     /label="Transposon right end"
     CDS             8679..9158
                     /label="ChlR    (Chloramphenicol    acetyltransferase)
(partial)"
     promoter        9159..9193
                     /label="J23119 promoter"
     CDS             9194..9373
                     /label="ChlR    (Chloramphenicol    acetyltransferase)
(partial)"
     misc_feature    complement(9410..9554)
                     /label="Transposon left end"
     misc_feature    complement(9547..9554)
                     /label="Transposon site duplication"
     CDS             complement(9887..10549)
                     /label="pBBR1 Rep"
     rep_origin      complement(10550..11321)
                     /label="pBBR1 oriV"
```

```
     misc_feature    11426..11477
                     /label="mob promoter and oriT"
     misc_feature    11545..12543
                     /label="mob gene"
     CDS             12801..13595
                     /label="NeoR/KanR          (Aminoglycoside          3'-
phosphotransferase)"
ORIGIN
        1 ttgacagcta gctcagtcct aggtataata ctagcgtcga cgtggagata taccatgggt
       61 gaactgccga gtaggtagct gataacgaga cctctgtctt gtcagctagg gtggtctcgt
      121 gaactgccga gtaggtagct gataacggat ccgaattcga gcgaaggaga tatacatatg
      181 tttttgcaaa gacctaaacc ttacagcgat gaaagtttag aaagtttctt tatccgagtg
      241 gctaacaaaa atggctacgg tgatgtccat cgcttcctag aagccactaa acgattcctt
      301 caagacattg accataatgg ctatcaaacc tttccgactg atataactcg gataaaccca
      361 tactcagcta aaaacagttc cagcgcacga actgcgtcat tcctgaaact tgcacaattg
      421 acatttaatg aaccgccaga gctacttggg ttggcaatta acagaacaaa catgaaatac
      481 tcgccgtcaa ctagcgcggt tgttagaggt gcagaagtct ttcctcgcag tttactacgg
      541 acgcactcca tcccctgctg tcctttgtgt ctgcgagaaa atggctacgc ctcctacctt
      601 tggcactttc aggggtacga atactgccac agccataacg tacctttaat taccacttgt
      661 agctgtggta aggagtttga ctaccgagta tctgggttaa agggcatttg ctgcaaatgc
      721 aaggagccta tcaccttaac cagcagggag aacggtcatg aggcagcgtg tactgtttca
      781 aactggcttg ctggccatga atctaaacct ctgccaaatc ttcctaaaag ctaccgatgg
      841 ggtttagttc attggtggat gggtattaaa gatagcgagt tcgatcactt ttcgttcgtt
      901 caatttttct caaactggcc aaggtcattc cactcgataa tcgaagatga agtagagttc
      961 aaccttgagc atgctgttgt cagcacgtct gaattacgac taaaagatct tcttggtcga
     1021 ttgtttttcg gttcaattcg gttacctgag cggaatcttc aacacaatat catccttggt
     1081 gagcttctct gctatttaga aaatcgttta tggcaagaca agggattaat cgccaacctc
     1141 aaaatgaacg cgttagaggc gactgtaatg ttaaattgta gcctcgatca gattgcatca
     1201 atggttgaac aacgcatctt gaagccaaat cgaaaaagca agcccaacag ccctcttgat
     1261 gttaccgatt atctatttca tttcggcgat attttctgtc tttggttagc tgagttccaa
     1321 agcgatgagt ttaaccgttc gttttatgtg tcgaggtggt aaatgcaaac tctgaaagaa
     1381 ctaatcgcat ccaatcctga cgacttaaca actgaactta agagagcatt tcgtccactc
     1441 acaccgcata ttgcaattga tggtaatgaa cttgacgcac tgacgatatt agtcaattta
     1501 accgataaga ctgatgatca gaaagacctg ctcgatcgag ccaaatgcaa gcaaaaactt
     1561 cgagatgaaa aatggtgggc tagctgcata aattgcgtta attacagaca aagccataac
     1621 ccaaaattcc cggatatacg ttctgaaggc gtgattcgaa cccaagccct gggtgaatta
     1681 ccgagtttcc tactctcgtc ctcaaaaatc ccaccatacc attggtcata tagccatgat
     1741 tcaaaatacg tcaacaaaag cgcattcctc accaatgagt tttgttggga tggtgagatc
     1801 tcatgtttgg gtgagcttct taaagatgca gatcacccac tttggaatac tttgaaaaag
     1861 ttaggttgtt ctcaaaaaac ttgcaaagca atggcaaaac aactagctga tattactctc
     1921 acgactatca atgtcacact cgcaccaaat tacctgactc aaatatctct ccccgatagt
     1981 gatacatcct acatttcact ctctcctgtt gcatcgctat cgatgcaaag ccactttcat
     2041 cagaggctcc aagatgaaaa tcggcatagc gcgataacgc ggtttagccg aactaccaac
     2101 atgggggtca ctgcgatgac atgtggcggc gcatttagaa tgttaaagtc tggcgctaag
     2161 ttttccagcc cccctcatca ccgattaaac agtaaacgaa gttggttgac gtcagagcat
     2221 gttcagtcat taaaacagta ccagcgcctc aacaaaagcc tcatacctga aaactctcgg
     2281 attgcactcc gtagaaaata caaaatcgag cttcaaaata tggtcagatc ttggtttgca
     2341 atgcaagacc ataccccttga ttcgaacata cttatccaac atttgaatca tgacctatct
     2401 tacttaggag ccacaaaacg ttttgcatac gatccagcga tgaccaagct ctttactgag
     2461 cttttgaaac gagagttatc aaattcaatc aataatggtg agcaacacac taatggatcg
     2521 tttttagtcc taccgaatat cagggtttgt ggcgcaacag ctttaagctc cccggtaacg
     2581 gtggggattc catcacttac agctttcttt ggcttcgttc acgcatttga acggaatata
     2641 aatcgcacca cctcatcgtt tcgtgttgaa tcctttgcga tatgcgtcca tcaactacat
     2701 gtcgagaagc gaggtttgac agcagagttt gtggaaaaag gcgacgggac tatatccgct
     2761 cccgcgaccc gggatgactg gcagtgtgat gtcgtattta gccttatttt gaacaccaac
     2821 tttgctcaac atattgacca agatacgtta gttacatcac taccaaagcg attggctcgg
     2881 ggttcagcaa aaattgcgat tgatgacttt aaacatatca actcattctc gacattagaa
```

2941 acagcgatcg aatctctgcc aatagaagct ggtaggtggt tatcacttta cgcacagtca
3001 aacaataatc taagtgatct attagcagcc atgacagagg accatcagct catggcaagc
3061 tgcgtcggtt accacttgtt agaagagccc aaagataaac caaactccct cagaggttac
3121 aaacacgcta tcgccgagtg catcattgga ctcattaact caatcacctt tagctcagag
3181 actgatccca acacaatctt ttggtcgcta aagaactatc aaaactacct agtggtacag
3241 ccaaggagta tcaacgatga aactaccgac aaatctagcc tatgagcgct ctatcgaccc
3301 atcagatgtc tgttttttttg tcgtctggcc cgatgataga aaaacacctt taacctacaa
3361 ttctcgtact ctgctcgggc aaatggaagc ggcatcatta gcctatgatg tctcaggtca
3421 accaataaaa agtgccaccg ctgaggcgtt agctcaaggg aaccctcatc aagttgattt
3481 ctgccacgtt ccatacggtg cgagtcatat tgaatgcagt ttctccgtct cgttttcttc
3541 tgaactacgt caaccatata agtgtaactc aagcaaagtt aaacaaacgc tagtgcaatt
3601 agtcgagctc tacgaaacga aaatcggctg gactgagcta gcaacccgat atttgatgaa
3661 catttgcaac ggtaaatggc tgtggaaaaa tacccgtaaa gcctattgct ggaacattgt
3721 acttacacct tggccttgga acggggaaaa ggttggattt gaagatattc gtactaacta
3781 cacctcacgg caagacttta aaaataataa aaattggtct gctatagttg aaatgatcaa
3841 aaccgcattt tctagtactg atgggctggc gatatttgaa gtcagggcca ccttgcactt
3901 gccaacgaat gctatggtgc ggccaagcca agttttcaca gaaaaagaaa gtggcagtaa
3961 aagtaaatct aaaactcaaa acagtcgagt ttttcagagt acaactattg atggtgaacg
4021 atcgccaata ctaggggcct ttaaaacggg agcagctatt gcaaccattg acgactggta
4081 tcctgaagcc actgagccac taagggtcgg acggtttggg gttcatcgcg aagatgtcac
4141 ttgctaccgt catccgtcta ccggaaaaga ttttttctcg atattacaac aagcagagca
4201 ctatattgaa gtgttgagcg ccaacaaaac tcccgctcaa gaaactatca acgacatgca
4261 ctttttaatg gctaacctga ttaagggtgg gatgttccag cataaaggag actgactgtg
4321 aaatggtatt ataagacaat cacctttctg ccagagttgt gcaacaacga gtcactggct
4381 gcaaagtgtc tccgcgttct gcatggattt aactatcagt atgagacacg aaatatcggc
4441 gtttcatttc cgctttggtg tgatgcaacg gttggaaaaa agatttcatt tgtcagcaag
4501 aacaagatag aactcgactt actacttaaa caacactatt tcgtccaaat ggaacaactt
4561 caatactttc atatatccaa cactgttctc gtcccagaag attgtacata cgtttccttt
4621 agacgctgtc aatctataga taagctcaca gcagcagggc tggcaaggaa aatcagacgc
4681 ctggagaaac gtgctctatc tagaggcgag caatttgacc catcatcttt tgctcaaaaa
4741 gagcatactg caatagcgca ctaccactca cttggggagt ccagcaaaca gacgaaccgc
4801 aactttcgac tcaatatcag gatgctctcg gagcaacccc gtgaggggaa ctcgattttt
4861 agtagctatg gcttatcaaa ttcagaaaac tcgtttcagc ctgtaccctt aatctgacct
4921 ttaataagga gatataccat ggcgacaagt ttacctacgc cctcagcaat tacgacttcg
4981 gcgttagagt atgcattcca tactcccgct cgcaatctaa cgaaatctcg cggaaaaaac
5041 attcatcgtt atgtcagtgt aaagatgagt aagaggatta cggtagaatc tactctagag
5101 tgtgatgcct gctatcactt tgattttgag ccaagtattg ttcgcttttg cgctcaaccg
5161 attcgatttt tatattatct caatggtcag tctcactcct atgttcctga ctttctagtt
5221 caatttgata ccaacgagtt tgttctatat gaagtaaagt cagcttatgc taagaacaaa
5281 cctgattttg atgttgaatg ggaggcgaaa gtaaaagcag caactgaact agggctagaa
5341 ttggagcttg ttgaagagag tgatattagg gatacggttg tattaaataa tcttaagcgc
5401 atgcatcgtt atgcttcgaa agatgagctg aataacgtac ataactctct cttaaaaata
5461 ataaagtaca atggcgccca atctgcaaga tgcttgggag aacagttggg tttaaaaggc
5521 cgaactgttt taccaatttt gtgcgatttg ctgtcaaggt gtttactcga tacacgtttg
5581 gataagcctc tatctcttga atctcgattt gagttggcca gttatggcta agaaagggtt
5641 ctcaagtttc catagaaaag cagtctcttc ccaagatacg cttgaatcga tagagcttgt
5701 ctctagcgct aattgtctag aaagtgttac gtatcaagat atatcagcat ttcccgaaac
5761 aattgcggta gagattaatt tccgattaag cattcttcgc tttttagcgc gaaagtgcga
5821 aaccattgtg gccaaatcaa ttgaaccaca tcgtgtagag ctacagcaaa actatagtag
5881 aaaaataccc agtgcaataa cgatatatcg atggtggctt gcttttcgaa aatcagacta
5941 caaccccatt agcttagcac ctaatatcaa ggatagaggt aatagagaga caaaagtgtc
6001 aacagttgtt gattctatta tggaacaggc agttgaaaga gttatatctg gacgaaaagt
6061 caatgttagc tctgcatata aacgtgttcg acgaaaagtt cgtcaataca atctaactca
6121 tggaacgaaa tacacgtatc ctaagtacga atctgtaaga aagcgagtaa aaaagaaaac
6181 cccatttgag ttattagccg cagggaaagg ggagagagta gctaagagag agtttcgccg
6241 aatgggaaaa aagatcctca cgtctagcgt gctagagagg gttgaaatag atcacactgt
6301 cgttgacctt tttgcagtac acgaagagta tcgaatccca ttgggccgac cttggcttac

6361 tcaattggtt gattgttaca gtaaagctgt aatcggtttt tatttaggtt tcgagcctcc
6421 tagctatgtg tcggtttccc ttgcacttaa gaatgcaata caacgcaaag atgacttaat
6481 ctcctcgtat gaatcgatcg agaatgaatg gctatgttat ggcatcccag acctactcgt
6541 aactgataat ggtaaagagt tttgtcgaa agcatttgat caagcatgtg aatcactatt
6601 gatcaatgtg catcaaaata aagttgagac gcccgacaac aaacctcatg ttgaacgtaa
6661 ctacgggact attaatactt ctctgttaga cgatttacct gggaaatcct tcagccagta
6721 ccttcaaaga gaagggtacg actctgtggg agaagctacc cttacactca atgagattag
6781 agaaatttac ttaatttggt tggtggatat ttatcataaa aaacccaatc agagaggcac
6841 taattgtcct aatgttgcgt ggaaaaaggg ttgtcaagaa tgggaaccag aggagttctc
6901 tggttctaaa gacgaattag actttaaatt tgctattgtt gattacaaac aacttactaa
6961 agtagggata actgtctaca aagaactgag ttatagcaat gaccgtttag ctgaatatag
7021 agggaagaaa ggaaaccata aagttcagtt caagtataac cctgagtgta tggcagttat
7081 ttgggtgttg gatgaggata tgaatgagta ctttacagtt aatgcgattg actacgaata
7141 tgcaagtaga gtatcacttt ggcaacataa atataacatg aaatatcaag cagaactaaa
7201 ttcagcagaa tatgatgagg acaaggaaat tgatgcagaa ataaaaattg aagaaatcgc
7261 agatcgttca attgttaaga ctaacaaaat cagagctcgg aggcgtggcg ctaggcatca
7321 agagaatagc gcaagggcta agtcaatcag taatgcgaac ccggcctcga tacaaaaaca
7381 tgaagatgaa atcgttagtg cagataatga cgattgggat attgattatg tctgagaaat
7441 cgtcaaatga gtgaaacgcg tgaggctcgt atatcaagag ctaaaagggc atttgtatcc
7501 acaccgagcg ttaggaaaat cttgagttac atggatagat gtagagatct atcagaccta
7561 gagtctgagc ctacatgcat gatggtctat ggtgcttcgg gtgtaggtaa aacgaccgtc
7621 atcaagaaat acttaaatca aaacagaaga gagtccgaag ccgggggcga tataataccg
7681 gttttgcata ttgagttgcc agacaatgcg aagccagtag atgcagcaag ggaattgctg
7741 gttgaaatgg gtgacccgct agcactttat gaaactgact tagctagatt gacgaaaaga
7801 ctgactgaat taatccctgc ggtcggcgtg aagctgatta ttatcgatga gttccaacat
7861 ttggtggaag aaaggtcaaa tcgggttctt acccaagtag gtaattggct aaaaatgata
7921 cttaacaaaa cgaaatgtcc aattgttata tttggtatgc catactcaaa agttgtactg
7981 caagcaaact cgcaacttca cgggcgattt tccattcagg ttgaactgcg ccccttttagc
8041 taccagggag gtagaggtgt atttaaaact tttttggaat accttgataa agccctacct
8101 tttgaaaaac aggctggctt agccaacgaa agtttgcaga aaaaattgta tgcattctct
8161 cagggaaaca tgcgttcgtt gagaaacctt atttatcaag catctatcga agcaattgat
8221 aatcagcatg agacgataac cgaagaagat ttcgtttttg catcgaagtt gacatcgggc
8281 gataaaccca actcatggaa aaatcctttt gaggagggtg ttgaggtaac agaagatatg
8341 ttacgaccgc caccaaaaga tattggttgg gaggactatt tgagacattc aaccccgaga
8401 gtgagtaaac caggtagaaa taaaaacttt ttcgaataac ctaggctgct gcgctgctgc
8461 caccgctgag caataactag cataacccct tggggcctct aaacgggtct tgaggggttt
8521 tttgctgaaa cctcaggcat ttgttgttga tacaaccata aaatgataat tacacccata
8581 aattgataat tatcacaccc ataaattgat attgcctctt catggtctaa acttcagtaa
8641 gtttacgaca ttttcctcga ggtcatttcc ggggatccat ggagaaaaaa atcactggat
8701 ataccaccgt tgatatatcc caatggcatc gtaaagaaca ttttgaggca tttcagtcag
8761 ttgctcaatg tacctataac cagaccgttc agctggatat tacggccttt ttaaagaccg
8821 taaagaaaaa taagcacaag ttttatccgg cctttattca cattcttgcc cgcctgatga
8881 atgctcatcc ggagttccgt atggcaatga aagacggtga gctggtgata tgggatagtg
8941 ttcacccttg ttacaccgtt ttccatgagc aaactgaaac gttttcatcg ctctggagtg
9001 aataccacga cgatttccgg cagtttctac acatatattc gcaagatgtg gcgtgttacg
9061 gtgaaaacct ggcctatttc cctaaagggt ttattgagaa tatgtttttc gtctcagcca
9121 atccctgggt gagtttcacc agttttgatt taaacgtgtt gacagctagc tcagtcctag
9181 gtataatact agcgccaata tggacaactt cttcgccccc gttttcacta tgggcaaata
9241 ttatacgcaa ggcgacaagg tgctgatgcc gctggcgatt caggttcatc atgccgtttg
9301 tgatggcttc catgtcggca gaatgcttaa tgaattacaa cagtactgcg atgagtggca
9361 gggcggggcg taattttttt aaggcagtta ttggtgccct tctagagtcc ttactgcagt
9421 agttttgctg aaatactcga ttcacaaaaa tatcaactta tggttgtttt gtgagatatc
9481 aatatatggt tgttttgtgg ttaagttgct gattataaat aattattaaa tatcacttta
9541 tggttgcatc aacaggtacc gagctcgaat tcactggccg tcgttttaca acgtcgtgac
9601 tgggaaaacc ctggcgttac ccaacttaat cgccttgcgg ccagtttgct caggctctcc
9661 ccgtggaggt aataattgac gatatgatca tttattctgc ctcccagagc ctgataaaaa
9721 cggtgaatcc gttagcgagg tgccgccggc ttccattcag gtcgaggtgg cccggctcca

9781 tgcaccgcga cgcaacgcgg ggaggcagac aaggtatagg gcggcgaggc ggctacagcc
9841 gatagtctgg aacagcgcac ttacgggttg ctgcgcaacc caagtgctac cggcgcggca
9901 gcgtgacccg tgtcggcggc tccaacggct cgccatcgtc cagaaaacac ggctcatcgg
9961 gcatcggcag gcgctgctgc ccgcgccgtt cccattcctc cgtttcggtc aaggctggca
10021 ggtctggttc catgcccgga atgccgggct ggctgggcgg ctcctcgccg gggccggtcg
10081 gtagttgctg ctcgcccgga tacagggtcg ggatgcggcg caggtcgcca tgccccaaca
10141 gcgattcgtc ctggtcgtcg tgatcaacca ccacggcggc actgaacacc gacaggcgca
10201 actggtcgcg gggctggccc cacgccacgc ggtcattgac cacgtaggcc gacacggtgc
10261 cggggccgtt gagcttcacg acggagatcc agcgctcggc caccaagtcc ttgactgcgt
10321 attggaccgt ccgcaaagaa cgtccgatga gcttggaaag tgtcttctgg ctgaccacca
10381 cggcgttctg gtggcccatc tgcgccacga ggtgatgcag cagcattgcc gccgtgggtt
10441 tcctcgcaat aagcccggcc cacgcctcat gcgctttgcg ttccgtttgc acccagtgac
10501 cgggcttgtt cttggcttga atgccgattt ctctggactg cgtggccatg cttatctcca
10561 tgcggtaggg gtgccgcacg gttgcggcac catgcgcaat cagctgcaac ttttcggcag
10621 cgcgacaaca attatgcgtt gcgtaaaagt ggcagtcaat tacagatttt ctttaaccta
10681 cgcaatgagc tattgcgggg ggtgccgcaa tgagctgttg cgtacccccc tttttaagt
10741 tgttgatttt taagtctttc gcatttcgcc ctatatctag ttctttggtg cccaaagaag
10801 ggcacccctg cggggttccc ccacgccttc ggcgcggctc cccctccggc aaaaagtggc
10861 ccctccgggg cttgttgatc gactgcgcgg ccttcggcct tgcccaaggt ggcgctgccc
10921 ccttggaacc cccgcactcg ccgccgtgag gctcgggggg caggcgggcg ggcttcgccc
10981 ttcgactgcc cccactcgca taggcttggg tcgttccagg cgcgtcaagg ccaagccgct
11041 gcgcggtcgc tgcgcgagcc ttgacccgcc ttccacttgg tgtccaaccg gcaagcgaag
11101 cgcgcaggcc gcaggccgga ggcttttccc cagagaaaat taaaaaaatt gatggggcaa
11161 ggccgcaggc cgcgcagttg gagccggtgg gtatgtggtc gaaggctggg tagccggtgg
11221 gcaatccctg tggtcaagct cgtgggcagg cgcagcctgt ccatcagctt gtccagcagg
11281 gttgtccacg ggccgagcga agcgagccag ccggtggccg ctcgcggcca tcgtccacat
11341 atccacgggc tggcaaggga gcgcagcgac cgcgcagggc gaagcccgga gagcaagccc
11401 gtagggcgcc gcagccgccg taggcggtca cgactttgcg aagcaaagtc tagtgagtat
11461 actcaagcat tgagtggccc gccggaggca ccgccttgcg ctgccccccgt cgagccggtt
11521 ggacaccaaa agggaggggc aggcatggcg gcatacgcga tcatgcgatg caagaagctg
11581 gcgaaaatgg gcaacgtggc ggccagtctc aagcacgcct accgcgagcg cgagacgccc
11641 aacgctgacg ccagcaggac gccagagaac gagcactggg cggccagcag caccgatgaa
11701 gcgatgggcc gactgcgcga gttgctgcca gagaagcggc gcaaggacgc tgtgttggcg
11761 gtcgagtacg tcatgacggc cagcccggaa tggtggaagt cggccagcca agaacagcag
11821 gcggcgttct tcgagaaggc gcacaagtgg ctggcggaca agtacggggc ggatcgcatc
11881 gtgacggcca gcatccaccg tgacgaaacc agcccgcaca tgaccgcgtt cgtggtgccg
11941 ctgacgcagg acggcaggct gtcggccaag gagttcatcg gcaacaaagc gcagatgacc
12001 cgcgaccaga ccacgtttgc ggccgctgtg gccgatctag ggctgcaacg ggcatcgag
12061 ggcagcaagg cacgtcacac gcgcattcag gcgttctacg aggccctgga gcggccacca
12121 gtgggccacg tcaccatcag cccgcaagcg gtcgagccac gcgcctatgc accgcaggga
12181 ttggccgaaa agctgggaat ctcaaagcgc gttgagacgc cggaagccgt ggccgaccgg
12241 ctgacaaaag cggttcggca ggggtatgag cctgccctac aggccgccgc aggagcgcgt
12301 gagatgcgca agaaggccga tcaagcccaa gagacggccc gtgaccttcg ggagcgcctg
12361 aagcccgttc tggacgccct ggggccgttg aatcgggata tgcaggccaa ggccgccgcg
12421 atcatcaagg ccgtgggcga aaagctgctg acggaacagc gggaagtcca gcgccagaaa
12481 caggcccagc gccagcagga acgcgggcgc gcacatttcc ccgaaaagtg ccacctggga
12541 tgaatgtcag ctactgggct atctggacaa gggaaaacgc aagcgcaaag agaaagcagg
12601 tagcttgcag tgggcttaca tggcgatagc tagactgggc ggttttatgg acagcaagcg
12661 aaccggaatt gccagctggg gcgccctctg gtaaggttgg gaagccctgc aaagtaaact
12721 ggatggcttt cttgccgcca aggatctgat ggcgcagggg atcaagatct gatcaagaga
12781 caggatgagg atcgtttcgc atgattgaac aagatggatt gcacgcaggt tctccggccg
12841 cttgggtgga gaggctattc ggctatgact gggcacaaca gacaatcggc tgctctgatg
12901 ccgccgtgtt ccggctgtca gcgcaggggc gcccggttct ttttgtcaag accgacctgt
12961 ccggtgccct gaatgaactg caagacgagg cagcgcggct atcgtggctg gccacgacgg
13021 gcgttccttg cgcagctgtg ctcgacgttg tcactgaagc gggaagggac tggctgctat
13081 tgggcgaagt gccggggcag gatctcctgt catctcacct tgctcctgcc gagaaagtat
13141 ccatcatggc tgatgcaatg cggcggctgc atacgcttga tccggctacc tgcccattcg

```
    13201 accaccaagc gaaacatcgc atcgagcgag cacgtactcg gatggaagcc ggtcttgtcg
    13261 atcaggatga tctggacgaa gagcatcagg ggctcgcgcc agccgaactg ttcgccaggc
    13321 tcaaggcgcg catgcccgac ggcgaggatc tcgtcgtgac ccatggcgat gcctgcttgc
    13381 cgaatatcat ggtggaaaat ggccgctttt ctggattcat cgactgtggc cggctgggtg
    13441 tggcggaccg ctatcaggac atagcgttgg ctacccgtga tattgctgaa gagcttggcg
    13501 gcgaatgggc tgaccgcttc ctcgtgcttt acggtatcgc cgctcccgat tcgcagcgca
    13561 tcgccttcta tcgccttctt gacgagttct tctgagcggg actctggggt tcgaaatgac
    13621 cgaccaagcg acgcccaacc tgccatcacg agatttcgat tccaccgccg ccttctatga
    13681 aaggttgggc ttcggaatcg ttttccggga cgccggctgg atgatcctcc agcgcgggga
    13741 tctcatgctg gagttcgcgt tggccgattc attaatgcag ctggcacgac aggtttcccg
    13801 actggtggat aaccgtatta ccgcctttga gtgagctgat accgctcgcc gcagccgaac
    13861 gaccgagcgc agcgagtcag tgagcgagga cacaagct
//
```

**sJEC042**

```
LOCUS       sJEC042          3972 bp ds-DNA     linear
FEATURES             Location/Qualifiers
     CDS             complement(1..507)
                     /label="lacZ (partial)"
     CDS             701..1504
                     /label="AprR   (Aminoglycoside   N(3)-acetyltransferase
IV)"
     terminator      1606..1658
                     /label="tVoigtS19, L3S1P00"
     terminator      complement(1659..1719)
                     /label="tVoigtS1, L3S2P21"
     misc_feature    1735..1755
                     /label="BBa_G00001"
     terminator      complement(1771..1800)
                     /label="t500"
     CDS             complement(1805..2521)
                     /label="SFGFP"
     misc_feature    complement(2522..2535)
                     /label="RBS"
     misc_feature    complement(2542..2582)
                     /label="HP14 Stability Hairpin"
     misc_feature    2610..2641
                     /label="mRFP1 crRNA"
     RBS             2710..2721
                     /label="BBa_B0034"
     misc_feature    2728..3405
                     /label="mRFP1"
     terminator      3410..3439
                     /label="t500"
     CDS             complement(3465..3972)
                     /label="lacZ (partial)"
ORIGIN
        1 catcatattt aatcagcgac tgatccaccc agtcccagac gaagccgccc tgtaaacggg
       61 gatactgacg aaacgcctgc cagtatttag cgaaaccgcc aagactgtta cccatcgcgt
      121 gggcgtattc gcaaaggatc agcgggcgcg tctctccagg tagcgaaagc catttttga
      181 tggaccattt cggcacagcc gggaagggct ggtcttcatc cacgcgcgcg tacatcgggc
      241 aaataatatc ggtggccgtg gtgtcggctc cgccgccttc atactgcacc gggcgggaag
      301 gatcgacaga tttgatccag cgatacagcg cgtcgtgatt agcgccgtgg cctgattcat
      361 tccccagcga ccagatgatc acactcgggt gattacgatc gcgctgcacc attcgcgtta
      421 cgcgttcgct catcgccggt agccagcgcg gatcatcggt cagacgattc attggcacca
      481 tgccgtgggt ttcaatattg gcttcattga tcttttctac ggggtctgac gctcagtgga
```

541 acgaaaactc acgttaaggg attttggtca tgagattatc aaaaaggatc ttcacctaga
601 tccttttggt tcatgtgcag ctccatcagc aaaaggggat gataagttta tcaccaccga
661 ctatttgcaa cagtgccgtt gatcgtgcta tgatcgactg atgtcatcag cggtggagtg
721 caatgtcgtg caatacgaat ggcgaaaagc cgagctcatc ggtcagcttc tcaaccttgg
781 ggttaccccc ggcggtgtgc tgctggtcca cagctccttc cgtagcgtcc ggcccctcga
841 agatgggcca cttggactga tcgaggccct gcgtgctgcg ctgggtccgg gagggacgct
901 cgtcatgccc tcgtggtcag gtctggacga cgagccgttc gatcctgcca cgtcgcccgt
961 tacaccggac cttggagttg tctctgacac attctggcgc ctgccaaatg taaagcgcag
1021 cgcccatcca tttgcctttg cggcagcggg gccacaggca gagcagatca tctctgatcc
1081 attgccccctg ccacctcact cgcctgcaag cccggtcgcc cgtgtccatg aactcgatgg
1141 gcaggtactt ctcctcggcg tgggacacga tgccaacacg acgctgcatc ttgccgagtt
1201 gatggcaaag gttccctatg ggtgccgag acactgcacc attcttcagg atggcaagtt
1261 ggtacgcgtc gattatctcg agaatgacca ctgctgtgag cgctttgcct tggcggacag
1321 gtggctcaag gagaagagcc ttcagaagga aggtccagtc ggtcatgcct ttgctcggtt
1381 gatccgctcc cgcgacattg tggcgacagc cctgggtcaa ctgggccgag atccgttgat
1441 cttcctgcat ccgccagagg cgggatgcga agaatgcgat gccgctcgcc agtcgattgg
1501 ctgagctcat gagcggagaa cgagatgacg ttggaggggc aaggtcgcgc tgattgctgg
1561 ggcaacacgt ggagcggatc ggggattgtc tttccaaata atgtagacga acaataaggg
1621 gagcgggaaa ccgctcccct tttttattga taacaaaagg accaaaacga aaaaaggccc
1681 ccctttcggg aggcctcttt tctggaattt ggtaccgagt taatcagtga ttaactgcag
1741 cggccgctac tagtacgtct cacagccagc aaaaaaaagc ccgcctttcg gcgggctttg
1801 gccattattt gtagagctca tccatgccat gtgtaatccc agcagcagtt acaaactcaa
1861 gaaggaccat gtggtcacgc ttttcgttgg gatctttcga aaggacagat tgtgtcgaca
1921 ggtaatggtt gtctggtaaa aggacagggc catcgccaat tggagtattt tgttgataat
1981 ggtctgctag ttgaacggaa ccatcttcaa cgttgtggcg aattttgaag ttagctttga
2041 ttccattctt ttgtttgtct gccgtgatgt atacattgtg tgagttaaag ttgtactcga
2101 gtttgtgtcc aagaatgttt ccatcttctt taaaatcaat accctttaac tcgatacgat
2161 taacaagggt atcaccttca aacttgactt cagcacgcgt cttgtaggtc ccgtcatctt
2221 tgaaagatat agtgcgttcc tgtacataac cttcgggcat ggcactcttg aaaaagtcat
2281 gccgtttcat gtgatccgga taacgggaaa agcattgaac accataggtc agagtagtga
2341 caagtgttgg ccacggaaca ggtagttttc cagtagtgca aataaattta agggtgagtt
2401 ttccgtttgt agcatcacct tcaccctctc cacggacaga aaatttgtgc ccattaacat
2461 caccatctaa ttcaacaaga attgggacaa ctccagtgaa aagttcttct cctttgctca
2521 tagatccttc ctcctagatc caaaatacgg taccgtcaac aatctcactc gagagtcgac
2581 gtatgtgcaa aactaagcat tccgaagcca ttgttagccg tatgaatagg gaaactaaac
2641 ccagtgataa gacctgatgt tttcgcttct ttaattacat ttggagattt tttatttaca
2701 gcactagtga aagaggagaa atactaaatg gcgagtagcg aagacgttat caaagagttc
2761 atgcgtttca aagttcgtat ggaaggttcc gttaacggtc acgagttcga aatcgaaggt
2821 gaaggtgaag gtcgtccgta cgaaggtacc cagaccgcta aactgaaagt taccaaaggt
2881 ggtccgctgc cgttcgcttg ggacatcctg tccccgcagt tccagtacgg ttccaaagct
2941 tacgttaaac acccggctga catcccggac tacctgaaac tgtccttccc ggaaggtttc
3001 aaatgggaac gtgttatgaa cttcgaagac ggtggtgttg ttaccgttac ccaggactcc
3061 tccctgcaag acggtgagtt catctacaaa gttaaactgc gtggtaccaa cttcccgtcc
3121 gacggtccgg ttatgcagaa aaaaaccatg ggttgggaag cttccaccga acgtatgtac
3181 ccggaagacg gtgctctgaa aggtgaaatc aaaatgcgtc tgaaactgaa agacggtggt
3241 cactacgacg ctgaagttaa aaccacctac atggctaaaa aaccggttca gctgccgggt
3301 gcttacaaaa ccgacatcaa actggacatc acctcccaca acgaagacta caccatcgtt
3361 gaacagtacg aacgtgctga aggtcgtcac tccaccggtg cttaatggcc aaagcccgcc
3421 gaaaggcggg cttttttttg ctggctgtga gacgtactag tacataaagt tgttctgctt
3481 catcagcagg atatcctgca ccatcgtctg ctcatccatg acctgaccat gcagaggatg
3541 atgctcgtga cggttaacgc ctcgaatcag caacggcttg ccgttcagca gcagcagacc
3601 attttcaatc cgcacctcgc ggaaaccgac atcgcaggct tctgcttcaa tcagcgtgcc
3661 gtcggcggtg tgcagttcaa ccaccgcacg atagagattc gggatttcgg cgctccacag
3721 tttcgggttt tcgacgttca gacgtagtgt gacgcgatcg gcataaccac cacgctcatc
3781 gataatttca ccgccgaaag gcgcggtgcc gctggcgacc tgcgtttcac cctgccataa
3841 agaaactgtt acccgtaggt agtcacgcaa ctcgccgcac atctgaactt cagcctccag
3901 tacagcgcgg ctgaaatcat cattaaagcg agtggcaaca tggaaatcgc tgatttgtgt

```
     3961 agtcggttta tg
//
```

**sJEC081**

```
LOCUS       sJEC081          5019 bp ds-DNA     linear
FEATURES             Location/Qualifiers
     CDS             complement(1..507)
                     /label="lacZ (partial)"
     CDS             701..1504
                     /label="AprR   (Aminoglycoside   N(3)-acetyltransferase
IV)"
     terminator      1606..1658
                     /label="tVoigtS19, L3S1P00"
     terminator      complement(1659..1719)
                     /label="tVoigtS1, L3S2P21"
     misc_feature    1735..1755
                     /label="BBa_G00001"
     terminator      complement(1771..1800)
                     /label="t500"
     CDS             complement(1805..2521)
                     /label="SFGFP"
     misc_feature    complement(2522..2535)
                     /label="RBS"
     misc_feature    complement(2542..2582)
                     /label="HP14 Stability Hairpin"
     misc_feature    2610..2641
                     /label="mRFP1 crRNA"
     misc_feature    2710..2717
                     /label="Transposon site duplication"
     misc_feature    2710..2820
                     /label="Transposon right end"
     CDS             2844..3323
                     /label="ChlR    (Chloramphenicol    acetyltransferase)
(partial)"
     promoter        3324..3358
                     /label="J23119"
     CDS             3359..3538
                     /label="ChlR    (Chloramphenicol    acetyltransferase)
(partial)"
     misc_feature    complement(3575..3719)
                     /label="Transposon left end"
     misc_feature    complement(3712..3719)
                     /label="Transposon site duplication"
     RBS             3757..3768
                     /label="BBa_B0034"
     misc_feature    3775..4452
                     /label="mRFP1"
     terminator      4457..4486
                     /label="t500"
     CDS             complement(4512..5019)
                     /label="lacZ (partial)"
ORIGIN
        1 catcatattt aatcagcgac tgatccaccc agtcccagac gaagccgccc tgtaaacggg
       61 gatactgacg aaacgcctgc cagtatttag cgaaaccgcc aagactgtta cccatcgcgt
      121 gggcgtattc gcaaaggatc agcgggcgcg tctctccagg tagcgaaagc catttttga
      181 tggaccattt cggcacagcc gggaagggct ggtcttcatc cacgcgcgcg tacatcgggc
      241 aaataatatc ggtggccgtg gtgtcggctc cgccgccttc atactgcacc gggcgggaag
```

```
 301 gatcgacaga tttgatccag cgatacagcg cgtcgtgatt agcgccgtgg cctgattcat
 361 tccccagcga ccagatgatc acactcgggt gattacgatc gcgctgcacc attcgcgtta
 421 cgcgttcgct catcgccggt agccagcgcg gatcatcggt cagacgattc attggcacca
 481 tgccgtgggt ttcaatattg gcttcattga tcttttctac ggggtctgac gctcagtgga
 541 acgaaaactc acgttaaggg attttggtca tgagattatc aaaaaggatc ttcacctaga
 601 tccttttggt tcatgtgcag ctccatcagc aaaaggggat gataagttta tcaccaccga
 661 ctatttgcaa cagtgccgtt gatcgtgcta tgatcgactg atgtcatcag cggtggagtg
 721 caatgtcgtg caatacgaat ggcgaaaagc cgagctcatc ggtcagcttc tcaaccttgg
 781 ggttaccccc ggcggtgtgc tgctggtcca cagctccttc cgtagcgtcc ggcccctcga
 841 agatgggcca cttggactga tcgaggccct gcgtgctgcg ctgggtccgg gagggacgct
 901 cgtcatgccc tcgtggtcag gtctggacga cgagccgttc gatcctgcca cgtcgcccgt
 961 tacaccggac cttggagttg tctctgacac attctggcgc ctgccaaatg taaagcgcag
1021 cgcccatcca tttgcctttg cggcagcggg gccacaggca gagcagatca tctctgatcc
1081 attgcccctg ccacctcact cgcctgcaag cccggtcgcc cgtgtccatg aactcgatgg
1141 gcaggtactt ctcctcggcg tgggacacga tgccaacacg acgctgcatc ttgccgagtt
1201 gatggcaaag gttccctatg ggtgccgag acactgcacc attcttcagg atggcaagtt
1261 ggtacgcgtc gattatctcg agaatgacca ctgctgtgag cgctttgcct tggcggacag
1321 gtggctcaag gagaagagcc ttcagaagga aggtccagtc ggtcatgcct ttgctcggtt
1381 gatccgctcc cgcgacattg tggcgacagc cctgggtcaa ctgggccgag atccgttgat
1441 cttcctgcat ccgccagagg cgggatgcga agaatgcgat gccgctcgcc agtcgattgg
1501 ctgagctcat gagcggagaa cgagatgacg ttggaggggc aaggtcgcgc tgattgctgg
1561 ggcaacacgt ggagcggatc ggggattgtc tttccaaata atgtagacga acaataaggg
1621 gagcgggaaa ccgctcccct tttttattga taacaaaagg accaaaacga aaaaaggccc
1681 ccctttcggg aggcctcttt tctggaattt ggtaccgagt taatcagtga ttaactgcag
1741 cggccgctac tagtacgtct cacagccagc aaaaaaaagc ccgcctttcg gcgggctttg
1801 gccattattt gtagagctca tccatgccat gtgtaatccc agcagcagtt acaaactcaa
1861 gaaggaccat gtggtcacgc ttttcgttgg gatctttcga aaggacagat tgtgtcgaca
1921 ggtaatggtt gtctggtaaa aggacagggc catcgccaat tggagtattt tgttgataat
1981 ggtctgctag ttgaacggaa ccatcttcaa cgttgtggcg aattttgaag ttagctttga
2041 ttccattctt ttgtttgtct gccgtgatgt atacattgtg tgagttaaag ttgtactcga
2101 gtttgtgtcc aagaatgttt ccatcttctt taaaatcaat accctttaac tcgatacgat
2161 taacaagggt atcaccttca aacttgactt cagcacgcgt cttgtaggtc ccgtcatctt
2221 tgaaagatat agtgcgttcc tgtacataac cttcgggcat ggcactcttg aaaaagtcat
2281 gccgtttcat gtgatccgga taacgggaaa agcattgaac accataggtc agagtagtga
2341 caagtgttgg ccacggaaca ggtagttttc cagtagtgca aataaattta agggtgagtt
2401 ttccgtttgt agcatcacct tcaccctctc cacggacaga aaatttgtgc ccattaacat
2461 caccatctaa ttcaacaaga attgggacaa ctccagtgaa aagttcttct cctttgctca
2521 tagatccttc ctcctagatc caaaatacgg taccgtcaac aatctcactc gagagtcgac
2581 gtatgtgcaa aactaagcat tccgaagcca ttgttagccg tatgaatagg gaaactaaac
2641 ccagtgataa gacctgatgt tttcgcttct ttaattacat ttggagattt tgaaacctca
2701 ggcatttgtt gttgatacaa ccataaaatg ataattacac ccataaattg ataattatca
2761 cacccataaa ttgatattgc ctcttcatgg tctaaacttc agtaagttta cgacattttc
2821 ctcgaggtca tttccgggga tccatggaga aaaaaatcac tggatatacc accgttgata
2881 tatcccaatg gcatcgtaaa gaacattttg aggcatttca gtcagttgct caatgtacct
2941 ataaccagac cgttcagctg gatattacgg cctttttaaa gaccgtaaag aaaaataagc
3001 acaagtttta tccggccttt attcacattc ttgcccgcct gatgaatgct catccggagt
3061 tccgtatggc aatgaaagac ggtgagctgg tgatatggga tagtgttcac ccttgttaca
3121 ccgttttcca tgagcaaact gaaacgtttt catcgctctg gagtgaatac cacgacgatt
3181 tccggcagtt tctacacata tattcgcaag atgtggcgtg ttacggtgaa aacctggcct
3241 atttccctaa agggtttatt gagaatatgt ttttcgtctc agccaatccc tgggtgagtt
3301 tcaccagttt tgatttaaac gtgttgacag ctagctcagt cctaggtata atactagcgc
3361 caatatggac aacttcttcg cccccgtttt cactatgggc aaatattata cgcaaggcga
3421 caaggtgctg atgccgctgg cgattcaggt tcatcatgcc gtttgtgatg gcttccatgt
3481 cggcagaatg cttaatgaat tacaacagta ctgcgatgag tggcagggcg gggcgtaatt
3541 tttttaaggc agttattggt gcccttctag agtccttact gcagtagttt tgctgaaata
3601 ctcgattcac aaaaatatca acttatggtt gttttgtgag atatcaatat atggttgttt
3661 tgtggttaag ttgctgatta taaataatta ttaaatatca ctttatggtt gcatcaacag
```

```
     3721 gtaccgagct cgaattcttt atttacagca ctagtgaaag aggagaaata ctaaatggcg
     3781 agtagcgaag acgttatcaa agagttcatg cgtttcaaag ttcgtatgga aggttccgtt
     3841 aacggtcacg agttcgaaat cgaaggtgaa ggtgaaggtc gtccgtacga aggtacccag
     3901 accgctaaac tgaaagttac caaaggtggt ccgctgccgt tcgcttggga catcctgtcc
     3961 ccgcagttcc agtacggttc caaagcttac gttaaacacc cggctgacat cccggactac
     4021 ctgaaactgt ccttcccgga aggtttcaaa tgggaacgtg ttatgaactt cgaagacggt
     4081 ggtgttgtta ccgttaccca ggactcctcc ctgcaagacg gtgagttcat ctacaaagtt
     4141 aaactgcgtg gtaccaactt cccgtccgac ggtccggtta tgcagaaaaa aaccatgggt
     4201 tgggaagctt ccaccgaacg tatgtacccg gaagacggtg ctctgaaagg tgaaatcaaa
     4261 atgcgtctga aactgaaaga cggtggtcac tacgacgctg aagttaaaac cacctacatg
     4321 gctaaaaaac cggttcagct gccgggtgct tacaaaaccg acatcaaact ggacatcacc
     4381 tcccacaacg aagactacac catcgttgaa cagtacgaac gtgctgaagg tcgtcactcc
     4441 accggtgctt aatggccaaa gcccgccgaa aggcgggctt ttttttgctg gctgtgagac
     4501 gtactagtac ataaagttgt tctgcttcat cagcaggata tcctgcacca tcgtctgctc
     4561 atccatgacc tgaccatgca gaggatgatg ctcgtgacgg ttaacgcctc gaatcagcaa
     4621 cggcttgccg ttcagcagca gcagaccatt ttcaatccgc acctcgcgga aaccgacatc
     4681 gcaggcttct gcttcaatca gcgtgccgtc ggcggtgtgc agttcaacca ccgcacgata
     4741 gagattcggg atttcggcgc tccacagttt cgggttttcg acgttcagac gtagtgtgac
     4801 gcgatcggca taaccaccac gctcatcgat aatttcaccg ccgaaaggcg cggtgccgct
     4861 ggcgacctgc gtttcaccct gccataaaga aactgttacc cgtaggtagt cacgcaactc
     4921 gccgcacatc tgaacttcag cctccagtac agcgcggctg aaatcatcat taaagcgagt
     4981 ggcaacatgg aaatcgctga tttgtgtagt cggtttatg
//
```

**sJEC050**

```
LOCUS       sJEC050          5019 bp ds-DNA     linear
FEATURES             Location/Qualifiers
     CDS             complement(1..507)
                     /label="lacZ (partial)"
     CDS             701..1504
                     /label="AprR   (Aminoglycoside   N(3)-acetyltransferase
IV)"
     terminator      1606..1658
                     /label="tVoigtS19, L3S1P00"
     terminator      complement(1659..1719)
                     /label="tVoigtS1, L3S2P21"
     misc_feature    1735..1755
                     /label="BBa_G00001"
     terminator      complement(1771..1800)
                     /label="t500"
     CDS             complement(1805..2521)
                     /label="SFGFP"
     misc_feature    complement(2522..2535)
                     /label="RBS"
     misc_feature    complement(2542..2582)
                     /label="HP14 Stability Hairpin"
     misc_feature    2610..2641
                     /label="mRFP1 crRNA"
     misc_feature    2709..2716
                     /label="Transposon site duplication"
     misc_feature    2709..2853
                     /label="Transposon left end"
     CDS             complement(2890..3069)
                     /label="ChlR    (Chloramphenicol    acetyltransferase)
(partial)"
     promoter        complement(3070..3104)
                     /label="J23119"
```

```
     CDS             complement(3105..3584)
                     /label="ChlR    (Chloramphenicol    acetyltransferase)
(partial)"
     misc_feature    complement(3608..3718)
                     /label="Transposon right end"
     misc_feature    complement(3711..3718)
                     /label="Transposon site duplication"
     RBS             3757..3768
                     /label="BBa_B0034"
     misc_feature    3775..4452
                     /label="mRFP1"
     terminator      4457..4486
                     /label="t500"
     CDS             complement(4512..5019)
                     /label="lacZ (partial)"
ORIGIN
        1 catcatattt aatcagcgac tgatccaccc agtcccagac gaagccgccc tgtaaacggg
       61 gatactgacg aaacgcctgc cagtatttag cgaaaccgcc aagactgtta cccatcgcgt
      121 gggcgtattc gcaaaggatc agcgggcgcg tctctccagg tagcgaaagc cattttttga
      181 tggaccattt cggcacagcc gggaagggct ggtcttcatc cacgcgcgcg tacatcgggc
      241 aaataatatc ggtggccgtg gtgtcggctc cgccgccttc atactgcacc gggcgggaag
      301 gatcgacaga tttgatccag cgatacagcg cgtcgtgatt agcgccgtgg cctgattcat
      361 tccccagcga ccagatgatc acactcgggt gattacgatc gcgctgcacc attcgcgtta
      421 cgcgttcgct catcgccggt agccagcgcg gatcatcggt cagacgattc attggcacca
      481 tgccgtgggt ttcaatattg gcttcattga tcttttctac ggggtctgac gctcagtgga
      541 acgaaaactc acgttaaggg attttggtca tgagattatc aaaaaggatc ttcacctaga
      601 tccttttggt tcatgtgcag ctccatcagc aaaaggggat gataagttta tcaccaccga
      661 ctatttgcaa cagtgccgtt gatcgtgcta tgatcgactg atgtcatcag cggtggagtg
      721 caatgtcgtg caatacgaat ggcgaaaagc cgagctcatc ggtcagcttc tcaaccttgg
      781 ggttaccccc ggcggtgtgc tgctggtcca cagctccttc cgtagcgtcc ggcccctcga
      841 agatgggcca cttggactga tcgaggccct gcgtgctgcg ctgggtccgg gagggacgct
      901 cgtcatgccc tcgtggtcag gtctggacga cgagccgttc gatcctgcca cgtcgcccgt
      961 tacaccggac cttggagttg tctctgacac attctggcgc ctgccaaatg taaagcgcag
     1021 cgcccatcca tttgcctttg cggcagcggg gccacaggca gagcagatca tctctgatcc
     1081 attgcccctg ccacctcact cgcctgcaag cccggtcgcc cgtgtccatg aactcgatgg
     1141 gcaggtactt ctcctcggcg tgggacacga tgccaacacg acgctgcatc ttgccgagtt
     1201 gatggcaaag gttccctatg ggtgccgag acactgcacc attcttcagg atggcaagtt
     1261 ggtacgcgtc gattatctcg agaatgacca ctgctgtgag cgctttgcct tggcggacag
     1321 gtggctcaag gagaagagcc ttcagaagga aggtccagtc ggtcatgcct ttgctcggtt
     1381 gatccgctcc cgcgacattg tggcgacagc cctgggtcaa ctgggccgag atccgttgat
     1441 cttcctgcat ccgccagagg cgggatgcga agaatgcgat gccgctcgcc agtcgattgg
     1501 ctgagctcat gagcggagaa cgagatgacg ttggaggggc aaggtcgcgc tgattgctgg
     1561 ggcaacacgt ggagcggatc ggggattgtc tttccaaata atgtagacga acaataaggg
     1621 gagcgggaaa ccgctcccct tttttattga taacaaaagg accaaaacga aaaaaggccc
     1681 ccctttcggg aggcctcttt tctggaattt ggtaccgagt taatcagtga ttaactgcag
     1741 cggccgctac tagtacgtct cacagccagc aaaaaaaagc ccgcctttcg gcgggctttg
     1801 gccattattt gtagagctca tccatgccat gtgtaatccc agcagcagtt acaaactcaa
     1861 gaaggaccat gtggtcacgc ttttcgttgg gatctttcga aaggacagat tgtgtcgaca
     1921 ggtaatggtt gtctggtaaa aggacagggc catcgccaat tggagtattt tgttgataat
     1981 ggtctgctag ttgaacggaa ccatcttcaa cgttgtggcg aattttgaag ttagctttga
     2041 ttccattctt ttgtttgtct gccgtgatgt atacattgtg tgagttaaag ttgtactcga
     2101 gtttgtgtcc aagaatgttt ccatcttctt taaaatcaat accctttaac tcgatacgat
     2161 taacaagggt atcaccttca aacttgactt cagcacgcgt cttgtaggtc ccgtcatctt
     2221 tgaaagatat agtgcgttcc tgtacataac cttcgggcat ggcactcttg aaaaagtcat
     2281 gccgtttcat gtgatccgga taacgggaaa agcattgaac accataggtc agagtagtga
     2341 caagtgttgg ccacggaaca ggtagttttc cagtagtgca aataaattta agggtgagtt
     2401 ttccgtttgt agcatcacct tcaccctctc cacggacaga aaatttgtgc ccattaacat
```

```
     2461 caccatctaa ttcaacaaga attgggacaa ctccagtgaa aagttcttct cctttgctca
     2521 tagatccttc ctcctagatc caaaatacgg taccgtcaac aatctcactc gagagtcgac
     2581 gtatgtgcaa aactaagcat tccgaagcca ttgttagccg tatgaatagg gaaactaaac
     2641 ccagtgataa gacctgatgt tttcgcttct ttaattacat ttggagattt gaattcgagc
     2701 tcggtacctg ttgatgcaac cataaagtga tatttaataa ttatttataa tcagcaactt
     2761 aaccacaaaa caaccatata ttgatatctc acaaaacaac cataagttga tatttttgtg
     2821 aatcgagtat ttcagcaaaa ctactgcagt aaggactcta gaagggcacc aataactgcc
     2881 ttaaaaaaat tacgccccgc cctgccactc atcgcagtac tgttgtaatt cattaagcat
     2941 tctgccgaca tggaagccat cacaaacggc atgatgaacc tgaatcgcca gcggcatcag
     3001 caccttgtcg ccttgcgtat aatatttgcc catagtgaaa acgggggcga agaagttgtc
     3061 catattggcg ctagtattat acctaggact gagctagctg tcaacacgtt taaatcaaaa
     3121 ctggtgaaac tcacccaggg attggctgag acgaaaaaca tattctcaat aaacccttta
     3181 gggaaatagg ccaggttttc accgtaacac gccacatctt gcgaatatat gtgtagaaac
     3241 tgccggaaat cgtcgtggta ttcactccag agcgatgaaa acgtttcagt ttgctcatgg
     3301 aaaacggtgt aacaagggtg aacactatcc catatcacca gctcaccgtc tttcattgcc
     3361 atacggaact ccggatgagc attcatcagg cgggcaagaa tgtgaataaa ggccggataa
     3421 aacttgtgct tatttttctt tacggtcttt aaaaaggccg taatatccag ctgaacggtc
     3481 tggttatagg tacattgagc aactgactga aatgcctcaa aatgttcttt acgatgccat
     3541 tgggatatat caacggtggt atatccagtg atttttttct ccatggatcc ccggaaatga
     3601 cctcgaggaa aatgtcgtaa acttactgaa gtttagacca tgaagaggca atatcaattt
     3661 atgggtgtga taattatcaa tttatgggtg taattatcat tttatggttg tatcaacaac
     3721 aaatgcctga ggtttcattt atttacagca ctagtgaaag aggagaaata ctaaatggcg
     3781 agtagcgaag acgttatcaa agagttcatg cgtttcaaag ttcgtatgga aggttccgtt
     3841 aacggtcacg agttcgaaat cgaaggtgaa ggtgaaggtc gtccgtacga aggtacccag
     3901 accgctaaac tgaaagttac caaaggtggt ccgctgccgt tcgcttggga catcctgtcc
     3961 ccgcagttcc agtacggttc caaagcttac gttaaacacc cggctgacat cccggactac
     4021 ctgaaactgt ccttcccgga aggtttcaaa tgggaacgtg ttatgaactt cgaagacggt
     4081 ggtgttgtta ccgttaccca ggactcctcc ctgcaagacg gtgagttcat ctacaaagtt
     4141 aaactgcgtg gtaccaactt cccgtccgac ggtccggtta tgcagaaaaa aaccatgggt
     4201 tgggaagctt ccaccgaacg tatgtacccg gaagacggtg ctctgaaagg tgaaatcaaa
     4261 atgcgtctga aactgaaaga cggtggtcac tacgacgctg aagttaaaac cacctacatg
     4321 gctaaaaaac cggttcagct gccgggtgct tacaaaaccg acatcaaact ggacatcacc
     4381 tcccacaacg aagactacac catcgttgaa cagtacgaac gtgctgaagg tcgtcactcc
     4441 accggtgctt aatggccaaa gcccgccgaa aggcgggctt tttttgctg gctgtgagac
     4501 gtactagtac ataaagttgt tctgcttcat cagcaggata tcctgcacca tcgtctgctc
     4561 atccatgacc tgaccatgca gaggatgatg ctcgtgacgg ttaacgcctc gaatcagcaa
     4621 cggcttgccg ttcagcagca gcagaccatt ttcaatccgc acctcgcgga aaccgacatc
     4681 gcaggcttct gcttcaatca gcgtgccgtc ggcggtgtgc agttcaacca ccgcacgata
     4741 gagattcggg atttcggcgc tccacagttt cgggttttcg acgttcagac gtagtgtgac
     4801 gcgatcggca taaccaccac gctcatcgat aatttcaccg ccgaaaggcg cggtgccgct
     4861 ggcgacctgc gtttcaccct gccataaaga aactgttacc cgtaggtagt cacgcaactc
     4921 gccgcacatc tgaacttcag cctccagtac agcgcggctg aaatcatcat taaagcgagt
     4981 ggcaacatgg aaatcgctga tttgtgtagt cggtttatg
//
```

**sJEC051**

```
LOCUS       sJEC051          3785 bp ds-DNA     linear
FEATURES             Location/Qualifiers
     gene            13..708
                     /label="yjaH"
     CDS             13..708

/translation="VNSFNEGVVSPLLSFWRRSLMLAGALLLTACSHNSSLPPFTASGFAEDQGAVRIWRKDSGDN
VHLLAVFSPWRSGDTTTREYRWQGDNLTLININVYSKPPVNIRARFDDRGDLSFMQRESDGEKQQLSNDQIDLYRY
RADQIRQISDALRQGRVVLRQGRWHAMEQTVTTCEGQTIKPDLDSQAIAHIERRQSRSSVDVSVAWLEAPEGSQLL
LVANSDFCRWQPNEKTF*"
     gene            complement(710..1135)
```

```
                     /label="zraP"
     CDS             complement(710..1135)

/translation="MKRNTKIALVMMALSAMAMGSTSAFAHGGHGMWQQNAAPLTSEQQTAWQKIHNDFYAQSSAL
QQQLVTKRYEYNALLAANPPDSSKINAVAKEMENLRQSLDELRVKRDIAMAEAGIPRGAGMGMGYGGCGGGGHMGM
GHW*"
     misc_feature    1223..1227
                     /label="Transposon site duplication"
     misc_feature    1228..1235
                     /label="Transposon site duplication"
     misc_feature    1228..1372
                     /label="Transposon left end"
     CDS             complement(1409..1588)
                     /label="ChlR    (Chloramphenicol    acetyltransferase)
(partial)"
     promoter        complement(1589..1623)
                     /label="J23119 promoter"
     CDS             complement(1624..2103)
                     /label="ChlR    (Chloramphenicol    acetyltransferase)
(partial)"
     misc_feature    complement(2127..2237)
                     /label="Transposon right end"
     misc_feature    complement(2230..2237)
                     /label="Transposon site duplication"
     misc_feature    2238..2242
                     /label="Transposon site duplication"
     misc_feature    complement(2289..2320)
                     /label="zraP crRNA"
     gene            2388..3785
                     /label="zraS"
     CDS             2388..3785

/translation="MRFMQRSKDSLAKWLSAILPVVIVGLVGLFAVTVIRDYGRASEADRQALLEKGNVLIRALES
GSRVGMGMRMHHVQQQALLEEMAGQPGVLWFAVTDAQGIIILHSDPDKVGRALYSPDEMQKLKPEENSRWRLLGKT
ETTPALEVYRLFQPMSAPWRHGMHNMPRCNGKAVPQVDAQQAIFIAVDASDLVATQSGEKRNTLIILFALATVLLA
SVLSFFWYRRYLRSRQLLQDEMKRKEKLVALGHLAAGVAHEIRNPLSSIKGLAKYFAERAPAGGEAHQLAQVMAKE
ADRLNRVVSELLELVKPTHLALQAVDLNTLINHSLQLVSQDANSREIQLRFTANDTLPEIQADPDRLTQVLLNLYL
NAIQAIGQHGVISVTASESGAGVKISVTDSGKGIAADQLDAIFTPYFTTKAEGTGLGLAVVHNIVEQHGGTIQVAS
QEGKGSTFTLWLPVNITRKDPQG*"
ORIGIN
        1 gattgcgtgg cagtgaacag ttttaacgaa ggggtggttt cacccctttt gtctttctgg
       61 cgtcgatcat tgatgctggc tggcgctctg cttctcactg cctgtagtca taactcttca
      121 cttcctccct ttaccgccag tggatttgct gaagaccagg gcgcggtacg catctggcga
      181 aaagacagcg gcgataatgt gcatctgctt gccgtgttta gcccgtggcg cagtggcgat
      241 accacgacgc gagagtatcg ctggcagggc gataacctca cgctcatcaa tatcaatgtt
      301 tacagcaaac cgccggtgaa tattcgcgcg cgttttgacg atcgcggtga tctgagcttt
      361 atgcaacgtg aatccgatgg ggaaaagcag cagctttcta acgaccaaat cgatttatac
      421 cgttatcgtg ctgatcagat ccgccagatt agcgatgcct tacgtcaggg gagagtcgtg
      481 ctgcgccagg ggcgctggca tgcgatggaa cagaccgtga ccacctgcga agggcaaacc
      541 attaaacctg atttagattc gcaggcgata gcgcatatcg agcgccgcca gagccgctct
      601 tctgttgatg tcagcgtggc atggctggaa gcgcccgaag gttcgcaatt actgttagtg
      661 gcaaactctg atttctgtcg ctggcaaccc aacgagaaaa cgttctgatt taccagtggc
      721 ccatacccat atgaccgcca ccaccgcagc cgccgtagcc catgcccatt ccggctccgc
      781 gcggaatacc cgcttcagcc atcgcgatat ctcgtttcac ccgtaactca tctaacgact
      841 gacgcaaatt ctccatctct ttggcgaccg cgttaatttt gctgctatcc ggtgggttcg
      901 cggctaacag ggcattgtat tcataacgct tcgtcaccag ttgctgttgc agtgcgctgc
      961 tttgagcgta aaagtcatta tggattttct gccacgctgt ctgttgttcg ctggtcaaag
```

```
     1021 gcgcggcatt ttgctgccac ataccgtgtc cgccgtgagc aaatgcagat gtcgatccca
     1081 tcgccattgc tgaaagcgcc atcattacca gggcaatttt cgtgttccgt ttcatggtta
     1141 atcctccagt ggttgtctta cttcgggtat tgcatcttcc gtgccaacga tgaaacgctg
     1201 atatgacggg taatctggca tgataaatgt tgatgcaacc ataaagtgat atttaataat
     1261 tatttataat cagcaactta accacaaaac aaccatatat tgatatctca caaaacaacc
     1321 ataagttgat atttttgtga atcgagtatt tcagcaaaac tactgcagta aggactctag
     1381 aagggcacca ataactgcct taaaaaaatt acgccccgcc ctgccactca tcgcagtact
     1441 gttgtaattc attaagcatt ctgccgacat ggaagccatc acaaacggca tgatgaacct
     1501 gaatcgccag cggcatcagc accttgtcgc cttgcgtata atatttgccc atagtgaaaa
     1561 cgggggcgaa gaagttgtcc atattggcgc tagtattata cctaggactg agctagctgt
     1621 caacacgttt aaatcaaaac tggtgaaact cacccaggga ttggctgaga cgaaaaacat
     1681 attctcaata aaccctttag ggaaataggc caggttttca ccgtaacacg ccacatcttg
     1741 cgaatatatg tgtagaaact gccggaaatc gtcgtggtat tcactccaga gcgatgaaaa
     1801 cgtttcagtt tgctcatgga aaacggtgta acaagggtga acactatccc atatcaccag
     1861 ctcaccgtct ttcattgcca tacggaactc cggatgagca ttcatcaggc gggcaagaat
     1921 gtgaataaag gccggataaa acttgtgctt atttttcttt acggtcttta aaaaggccgt
     1981 aatatccagc tgaacggtct ggttataggt acattgagca actgactgaa atgcctcaaa
     2041 atgttcttta cgatgccatt gggatatatc aacggtggta tatccagtga tttttttctc
     2101 catggatccc cggaaatgac ctcgaggaaa atgtcgtaaa cttactgaag tttagaccat
     2161 gaagaggcaa tatcaattta tgggtgtgat aattatcaat ttatgggtgt aattatcatt
     2221 ttatggttgt atcaacaata aacgagtaaa aatgactcgc ctgctgcggg tagcgagtca
     2281 tttttactca ttgaaactta agcctttgtg ttacagcgca gggtaagcgc tgataaaaga
     2341 tggcatgatt tctgctgtca gaaagggatg agcaggcaaa gaagaagatg cgttttatgc
     2401 aacgttctaa agactcctta gctaaatggt taagcgcgat cctccccgtg gtcattgttg
     2461 ggctggtggg attgtttgcg gtaactgtga ttcgtgatta tggcgggca agcgaggcag
     2521 accgccaggc attactggaa aaaggtaatg tgcttatccg cgctctggag tcgggaagcc
     2581 gcgtagggat ggggatgcga atgcaccatg tacagcaaca ggcgcttctg gaagagatgg
     2641 cgggacagcc gggagtgttg tggttcgcag tcaccgatgc gcagggcatc attattcttc
     2701 atagcgaccc cgataaggtc gggcgtgcgc tctattcgcc ggatgaaatg cagaaattaa
     2761 agccagagga aaactcccgc tggcggctgc ttgggaaaac ggaaactacg cctgcacttg
     2821 aggtctatcg tttgttccag ccaatgtcag cgccctggcg gcatggaatg cacaatatgc
     2881 cgcgctgtaa cggcaaagct gtgccacaag tagatgcaca acaggctatt tttatcgccg
     2941 ttgatgccag tgatctggtt gcaacccaga gtggggaaaa gcgcaatacc ctgattatcc
     3001 tcttcgccct ggcgacggtc ttgctggcaa gcgtattgtc attcttctgg tatcgccgct
     3061 atctgcgctc gcgccagctt ctacaagatg aaatgaagcg caaagagaag ctggtggcgc
     3121 tggggcatct tgcggcaggc gttgcccacg aaatccgtaa cccactttcc tcgattaaag
     3181 gactggcgaa atactttgcc gagcgcgcgc ctgcaggggg agaagcgcat caactggcgc
     3241 aggtgatggc gaaagaggcc gaccgtttaa accgcgtggt aagcgagttg ctggaactgg
     3301 ttaagccaac gcatctggct ttgcaggcgg tggatctcaa cacgctgatt aaccactcat
     3361 tacagctggt aagtcaggat gcaaacagcc gggagatcca gttacgcttt accgccaacg
     3421 acacattacc ggaaattcag gccgacccgg acaggctgac tcaggtcctg ttgaatctct
     3481 atctcaatgc tattcaggcg attggtcagc atggcgtgat tagcgtgacg gccagcgaaa
     3541 gcggcgcggg cgtgaaaatc agcgttaccg acagcggtaa gggaattgcg gcagatcagc
     3601 ttgatgccat cttcactccg tacttcacca ctaaagccga aggcaccgga ttggggctgg
     3661 cggtcgtgca taatattgtt gaacaacacg gtggtacaat tcaggtcgca agccaggagg
     3721 gaaaaggctc aacgttcacc ctctggcttc cggtcaatat tacgcgtaag gacccacaag
     3781 gatga
//
```

**sJEC052**

```
LOCUS       sJEC052          4332 bp ds-DNA     linear
FEATURES             Location/Qualifiers
     gene            complement(80..1414)
                     /label="cadB"
     CDS             complement(80..1414)

/translation="MSSAKKIGLFACTGVVAGNMMGSGIALLPANLASIGGIAIWGWIISIIGAMSLAYVYARLAT
```

```
KNPQQGGPIAYAGEISPAFGFQTGVLYYHANWIGNLAIGITAVSYLSTFFPVLNDPVPAGIACIAIVWVFTFVNML
GGTWVSRLTTIGLVLVLIPVVMTAIVGWHWFDAATYAANWNTADTTDGHAIIKSILLCLWAFVGVESAAVSTGMVK
NPKRTVPLATMLGTGLAGIVYIAATQVLSGMYPSSVMAASGAPFAISASTILGNWAAPLVSAFTAFACLTSLGSWM
MLVGQAGVRAANDGNFPKVYGEVDSNGIPKKGLLLAAVKMTALMILITLMNSAGGKASDLFGELTGIAVLLTMLPY
FYSCVDLIRFEGVNIRNFVSLICSVLGCVFCFIALMGASSFELAGTFIVSLIILMFYARKMHERQSHSMDNHTASN
AH*"
     misc_feature    1493..1497
                     /label="Transposon site duplication"
     misc_feature    1498..1505
                     /label="Transposon site duplication"
     misc_feature    1498..1642
                     /label="Transposon left end"
     CDS             complement(1679..1858)
                     /label="ChlR    (Chloramphenicol    acetyltransferase)
(partial)"
     promoter        complement(1859..1893)
                     /label="J23119 promoter"
     CDS             complement(1894..2373)
                     /label="ChlR    (Chloramphenicol    acetyltransferase)
(partial)"
     misc_feature    complement(2397..2507)
                     /label="Transposon right end"
     misc_feature    complement(2500..2507)
                     /label="Transposon site duplication"
     misc_feature    2508..2512
                     /label="Transposon site duplication"
     misc_feature    complement(2556..2587)
                     /label="cadB crRNA"
     gene            complement(2794..4332)
                     /label="cadC"
     CDS             complement(2794..4332)

/translation="MQQPVVRVGEWLVTPSINQISRNGRQLTLEPRLIDLLVFFAQHSGEVLSRDELIDNVWKRSI
VTNHVVTQSISELRKSLKDNDEDSPVYIATVPKRGYKLMVPVIWYSEEEGEEIMLSSPPPIPEAVPATDSPSHSLN
IQNTATPPEQSPVKSKRFTTFWVWFFFLLSLGICVALVAFSSLDTRLPMSKSRILLNPRDIDINMVNKSCNSWSSP
YQLSYAIGVGDLVATSLNTFSTFMVHDKINYNIDEPSSSGKTLSIAFVNQRQYRAQQCFMSIKLVDNADGSTMLDK
RYVITNGNQLAIQNDLLESLSKALNQPWPQRMQETLQKILPHRGALLTNFYQAHDYLLHGDDKSLNRASELLGEIV
QSSPEFTYARAEKALVDIVRHSQHPLDEKQLAALNTEIDNIVTLPELNNLSIIYQIKAVSALVKGKTDESYQAINT
GIDLEMSWLNYVLLGKVYEMKGMNREAADAYLTAFNLRPGANTLYWIENGIFQTSVPYVVPYLDKFLASE*"
ORIGIN
        1 agtcatatct ccaggtaaaa aaggcccctc ccaacacatg ggacaaaatg aaaggaggag
       61 cctcggaaaa tacttttaat taatgtgcgt tagacgcggt gtggttatcc attgagtggc
      121 tctggcgctc gtgcattttg cgagcgtaga acatcaggat aatcaggctg acgatgaagg
      181 tacctgccag ctcgaaggag cttgcgccca tcagcgcgat gaagcagaac acgcaaccca
      241 gtacagagca gatcaggctg acaaagttgc ggatgttaac gccttcaaaa cgaatcaggt
      301 caacgcaaga gtagaaatac ggcagcatag tcagcagtac tgcgataccg gtcagttcac
      361 cgaacaggtc agatgcttta ccaccggcag agttcatcag agtgataagg atcatcaggg
      421 cagtcatttt cactgcagcc agcagcagac ctttttttcgg aataccgttg ctgtcgactt
      481 caccataaac tttcgggaag ttaccgtcgt tagcggcacg tacacctgcc tggcctacca
      541 acatcatcca ggagcccaga gaagtcaggc acgcaaaggc ggtgaatgca gaaaccagcg
      601 gcgcagccca gttaccgagg atagttgaag cactgattgc aaacggagca ccggaagccg
      661 ccattacaga agacggatac ataccggaaa gcacctgagt cgcagcgatg taaacaatac
      721 ctgctaaacc agtacccagc atggttgcca gcggaacggt acgtttcggg tttttaacca
      781 taccagtact tacagctgcg gattcaacac ccacgaaggc ccacaggcag agcagaatac
      841 ttttaatgat cgcatgacca tcagtggtat ccgcagtatt ccagttagct gcataagttg
      901 ccgcatcaaa ccaatgccag ccaacaatag cagtcatcac cacaggaata agaaccagca
      961 ccagaccaat agtggttaaa cggcttaccc aagtaccgcc gagcatattt acaaaggtaa
```

```
     1021 atacccagac gatagcaata caggcgatac ccgccggaac aggatcattt aatactggga
     1081 agaaggtgga aagataagat acagcggtaa taccaatcgc caggttacca atccagttag
     1141 catggtaata aagaacacct gtctgaaaac caaatgcagg ggaaatttct ccggcataag
     1201 caattgggcc accttgttgc gggttttttg ttgccagtcg ggcatataca tacgccagcg
     1261 acattgcacc aataatagag ataatccaac cccagatagc aataccaccg atacttgcta
     1321 ggttcgcagg taataatgca ataccgctcc ccatcatatt accggcaaca acaccggtac
     1381 aggcaaatag cccgatcttc ttggcagaac tcatgctctt ctcctaattt catttttgaa
     1441 tttggagtcc gggtcatgat gtataactat ttcctgacca gaccaaactg gcgataatgt
     1501 tgatgcaacc ataaagtgat atttaataat tatttataat cagcaactta accacaaaac
     1561 aaccatatat tgatatctca caaaacaacc ataagttgat atttttgtga atcgagtatt
     1621 tcagcaaaac tactgcagta aggactctag aagggcacca ataactgcct taaaaaaatt
     1681 acgccccgcc ctgccactca tcgcagtact gttgtaattc attaagcatt ctgccgacat
     1741 ggaagccatc acaaacggca tgatgaacct gaatcgccag cggcatcagc accttgtcgc
     1801 cttgcgtata atatttgccc atagtgaaaa cgggggcgaa gaagttgtcc atattggcgc
     1861 tagtattata cctaggactg agctagctgt caacacgttt aaatcaaaac tggtgaaact
     1921 cacccaggga ttggctgaga cgaaaaacat attctcaata aaccctttag ggaaataggc
     1981 caggttttca ccgtaacacg ccacatcttg cgaatatatg tgtagaaact gccggaaatc
     2041 gtcgtggtat tcactccaga gcgatgaaaa cgtttcagtt tgctcatgga aaacggtgta
     2101 acaagggtga acactatccc atatcaccag ctcaccgtct ttcattgcca tacggaactc
     2161 cggatgagca ttcatcaggc gggcaagaat gtgaataaag gccggataaa acttgtgctt
     2221 atttttcttt acggtcttta aaaaggccgt aatatccagc tgaacggtct ggttataggt
     2281 acattgagca actgactgaa atgcctcaaa atgttcttta cgatgccatt gggatatatc
     2341 aacggtggta tatccagtga tttttttctc catggatccc cggaaatgac ctcgaggaaa
     2401 atgtcgtaaa cttactgaag tttagaccat gaagaggcaa tatcaattta tgggtgtgat
     2461 aattatcaat ttatgggtgt aattatcatt ttatggttgt atcaacagat aagattactc
     2521 acgaaaaaag gattaatcct aaagattagg tgaaataaca caaaagtttc tgtaagtgag
     2581 aacttgaggt tttttattaa cacatcagga tcgcaagttg atatcatgaa aagataaaca
     2641 tttaatgttt acaatggatt gcgtgacatt ctctggttaa atttatgtaa taaaaattat
     2701 gcggcaaata aattgccgca acatattata ccaacaggaa catacaaaaa ctcaacaaca
     2761 aatatttccg agcataaatc aacccggagt tacttattct gaagcaagaa atttgtcgag
     2821 ataaggtaca acataaggaa cagaagtctg gaatatacca ttttcaatcc agtaaagggt
     2881 gtttgcccct gggcgtaaat taaaggcggt gagatatgca tcagctgctt cccggttcat
     2941 ccccttcatt tcataaacct tgccaagcaa cacataattt agccaggaca tttcaagatc
     3001 aatgccagta tttatcgcct ggtaagactc atctgtttta ccttttacca gagcactgac
     3061 cgcttttatt tgatatataa tggacaggtt gttcaattcc ggcagtgtaa caatgttatc
     3121 tatttctgtg ttcagtgctg ctaattgttt ttcatctaaa ggatgttgag aatggcgcac
     3181 gatatcaact aatgcttttt ctgctctcgc gtaggtaaat tctggggatg attgaacaat
     3241 ctcacctaat aattcactgg cacggttcaa tgatttatca tcgccatgca gtaaataatc
     3301 atgtgcctga taaaaattag ttaataacgc accacgatgc ggcaaaattt tctggagcgt
     3361 ctcctgcatt cgttgtggcc acggttggtt taacgctttt gataaactct ccagtaaatc
     3421 attttgaatc gccagctgat taccgttagt gatgacataa cgtttatcca gcatggttga
     3481 accatctgca ttgtctacca attttatcga cataaagcat tgttgagcac ggtattggcg
     3541 ctgattaaca aacgcaatag ataatgtttt accggaactg ctcggttcat caatgttgta
     3601 gttgattttg tcatgcacca taaaggtgga gaaggtgtta agtgatgtcg ccaccaaatc
     3661 acccacgcct atcgcgtaag agagctgata cggggaactc cagctgttac aactttttatt
     3721 taccatatta atgtcaatat cgcgtggatt gagcaaaata cgcgatttgc tcataggaag
     3781 acgtgtatca agacttgaaa acgctaccag tgctacacag atacctaacg acaacaggaa
     3841 aaaaaaccat acccaaaagg tagtgaatcg tttgctttta actggggatt gttcaggtgg
     3901 cgttgcggtg ttttgaatgt taagactgtg ggagggagaa tctgtggcag gaaccgcctc
     3961 tggtataggg ggaggcgaag atagcattat ttcctctccc tcttcttcgc tgtaccagat
     4021 aaccggcacc attaatttat agccgcgctt tggtacagta gcgatataga caggactatc
     4081 ttcatcatta tcttttaatg acttacgtag ttctgagata ctctgcgtca caacgtgatt
     4141 ggtgacaata cttctcttcc agacattatc gataagttca tccctgctaa gtacttcgcc
     4201 actgtgttga gcaaagaaaa ccagaagatc gattaatctc ggctcaaggg taagttgacg
     4261 cccattgcgg ctaatttggt ttatggacgg agtaacaagc cattcgccaa cgcgaactac
     4321 aggttgttgc at
//
```

**sJEC053**

```
LOCUS       sJEC053          3722 bp ds-DNA     linear
FEATURES             Location/Qualifiers
     gene            1..1215
                     /label="ydiM"
     CDS             1..1215

/translation="MKNPYFPTALGLYFNYLVHGMGVLLMSLNMASLETLWQTNAAGVSIVISSLGIGRLSVLLFA
GLLSDRFGRRPFIMLGMCCYMAFFFGILQTNNIIIAYVFGFLAGMANSFLDAGTYPSLMEAFPRSPGTANILIKAF
VSSGQFLLPLIISLLVWAELWFGWSFMIAAGIMFINALFLYRCTFPPHPGRRLPVIKKTTSSTEHRCSIIDLASYT
LYGYISMATFYLVSQWLAQYGQFVAGMSYTMSIKLLSIYTVGSLLCVFITAPLIRNTVRPTTLLMLYTFISFIALF
TVCLHPTFYVVIIFAFVIGFTSAGGVVQIGLTLMAERFPYAKGKATGIYYSAGSIATFTIPLITAHLSQRSIADIM
WFDTAIAAIGFLLALFIGLRSRKKTRHHSLKENVAPGG*"
     misc_feature    1277..1308
                     /label="ydiN crRNA"
     misc_feature    1353..1357
                     /label="Transposon site duplication"
     misc_feature    1358..1365
                     /label="Transposon site duplication"
     misc_feature    1358..1468
                     /label="Transposon right end"
     CDS             1492..1971
                     /label="ChlR    (Chloramphenicol    acetyltransferase)
(partial)"
     promoter        1972..2006
                     /label="J23119 promoter"
     CDS             2007..2186
                     /label="ChlR    (Chloramphenicol    acetyltransferase)
(partial)"
     misc_feature    complement(2223..2367)
                     /label="Transposon left end"
     misc_feature    complement(2360..2367)
                     /label="Transposon site duplication"
     misc_feature    2368..2372
                     /label="Transposon site duplication"
     gene            2457..3722
                     /label="ydiN"
     CDS             2457..3722

/translation="MSQNKAFSTPFILAVLCIYFSYFLHGISVITLAQNMSSLAEKFSTDNAGIAYLISGIGLGRL
ISILFFGVISDKFGRRAVILMAVIMYLLFFFGIPACPNLTLAYGLAVCVGIANSALDTGGYPALMECFPKASGSAV
ILVKAMVSFGQMFYPMLVSYMLLNNIWYGYGLIIPGILFVLITLMLLKSKFPSQLVDASVTNELPQMNSKPLVWLE
GVSSVLFGVAAFSTFYVIVVWMPKYAMAFAGMSEAEALKTISYYSMGSLVCVFIFAALLKKMVRPIWANVFNSALA
TITAAIIYLYPSPLVCNAGAFVIGFSAAGGILQLGVSVMSEFFPKSKAKVTSIYMMMGGLANFVIPLITGYLSNIG
LQYIIVLDFTFALLALITAIIVFIRYYRVFIIPENDVRFGERKFCTRLNTIKHRG*"
ORIGIN
        1 atgaaaaatc cctatttccc taccgcactt gggttgtatt ttaattacct ggtgcatggt
       61 atgggcgtcc ttttgatgag cctgaatatg gcctcgctgg agacactttg gcagactaat
      121 gccgcgggtg tctcgatagt tatctcatcg ctgggcattg gtcgattaag tgtcttgctt
      181 tttgcaggat tattatccga tcgctttggt cgccgccctt ttatcatgct cgggatgtgc
      241 tgctatatgg ccttcttttt tggcatcctg cagaccaata acatcattat cgcttatgtt
      301 tttggctttc tggcgggaat ggcaaacagt tttctcgatg caggcactta tcccagtttg
      361 atggaagctt ttccacgctc acctgggaca gccaatattt taattaaagc atttgtttcc
      421 agcggacaat tttattacc gctaatcatt agcctgttag tgtgggctga actgtggttc
      481 ggttggtcct ttatgattgc tgcaggcatt atgtttatta acgctctgtt tttataccgt
      541 tgtacgttcc caccccatcc gggtcgtcgc ttacctgtca taaagaaaac caccagctct
```

```
      601 acggaacatc gctgttcaat tatcgattta gccagttata ccttatatgg ctatatctca
      661 atggcaacgt tttatctggt tagccagtgg ctggcacagt acggacaatt tgttgcaggc
      721 atgtcataca ctatgtcgat caaactactc agtatctaca ccgtgggttc gctgctttgt
      781 gtatttatta ccgctccact cattcgtaat accgttcgcc caacaacatt actgatgctg
      841 tacaccttta tctcatttat tgctctgttt accgtctgcc tgcatcccac attttatgtg
      901 gtgataatat ttgcttttgt cattggtttt acctctgctg gaggtgttgt gcaaattggc
      961 ctgacgttaa tggctgaacg tttcccttac gctaaaggta aagctacagg gatctattac
     1021 agtgcgggca gtattgcgac ctttactatt ccgttgatta cggctcatct gtcccaaaga
     1081 agtattgccg atattatgtg gttcgatacc gccatcgctg ccatcggttt tttactggca
     1141 ctgtttatcg gcttacgcag ccgcaaaaaa acgcggcatc actcgctaaa ggaaaatgtc
     1201 gctccgggtg ggtaatgcaa tattcttttc aggtcatgca agatcttacg gataaataac
     1261 tctttctgcg ctaactaagg aaaatcgcga tcaaaaacaa actatgacat gcaatattcc
     1321 tggaaacata aactttatgc catgtaccca gggaaaatgt tgatacaacc ataaaatgat
     1381 aattacaccc ataaattgat aattatcaca cccataaatt gatattgcct cttcatggtc
     1441 taaacttcag taagtttacg acattttcct cgaggtcatt tccggggatc catggagaaa
     1501 aaaatcactg gatataccac cgttgatata tcccaatggc atcgtaaaga acattttgag
     1561 gcatttcagt cagttgctca atgtacctat aaccagaccg ttcagctgga tattacggcc
     1621 tttttaaaga ccgtaaagaa aaataagcac aagttttatc cggcctttat tcacattctt
     1681 gcccgcctga tgaatgctca tccggagttc cgtatggcaa tgaaagacgg tgagctggtg
     1741 atatgggata gtgttcaccc ttgttacacc gttttccatg agcaaactga aacgttttca
     1801 tcgctctgga gtgaatacca cgacgatttc cggcagtttc tacacatata ttcgcaagat
     1861 gtggcgtgtt acggtgaaaa cctggcctat ttccctaaag ggtttattga gaatatgttt
     1921 ttcgtctcag ccaatccctg ggtgagtttc accagttttg atttaaacgt gttgacagct
     1981 agctcagtcc taggtataat actagcgcca atatggacaa cttcttcgcc cccgttttca
     2041 ctatgggcaa atattatacg caaggcgaca aggtgctgat gccgctggcg attcaggttc
     2101 atcatgccgt ttgtgatggc ttccatgtcg gcagaatgct taatgaatta caacagtact
     2161 gcgatgagtg gcagggcggg gcgtaatttt tttaaggcag ttattggtgc ccttctagag
     2221 tccttactgc agtagttttg ctgaaatact cgattcacaa aaatatcaac ttatggttgt
     2281 tttgtgagat atcaatatat ggttgttttg tggttaagtt gctgattata aataattatt
     2341 aaatatcact ttatggttgc atcaacagaa aatcatcttc agtatagtaa ttatgtaaac
     2401 cgtcggagaa caatacgtac ggtaacgaaa ttatctttca gcaaggagct gtgaaaatgt
     2461 ctcaaaataa ggctttcagc acgccattta tcctggctgt tctttgtatt tacttcagct
     2521 acttcctgca cggcattagt gttattacgc ttgcccaaaa tatgtcatct ctggcggaaa
     2581 agttttccac tgacaacgcg ggcattgcct acttaatttc cggtatcggt ttggggcgat
     2641 tgatcagtat tttattcttc ggtgtgatct ccgataagtt tggtcgtcgg gcggtgatat
     2701 taatggcagt aataatgtat ctgctattct tctttggtat tcccgcttgc ccgaatttaa
     2761 ctctcgccta cggtctggca gtgtgcgtag gtatcgctaa ctcagcgctg gatacgggtg
     2821 gctaccccgc gctcatggaa tgctttccga aagcctctgg ttcggcggtc atactggtta
     2881 aagcgatggt gtcatttggg caaatgttct acccaatgct ggtgagctat atgttgctca
     2941 ataatatctg gtacggctat gggctgatta ttccgggtat tctatttgta ctgatcacgc
     3001 tgatgctgtt gaaaagcaaa ttccccagcc agttggtgga cgccagcgta actaatgaat
     3061 taccgcaaat gaacagcaaa ccgttagtct ggctggaagg tgtttcatcg gtactgttcg
     3121 gtgtagccgc attctcgacc ttttatgtga ttgtggtgtg gatgcccaaa tatgcgatgg
     3181 cttttgctgg tatgtcagaa gctgaggcat taaaaaccat ctcttattac agtatgggct
     3241 cgttggtctg tgtctttatt tttgccgcac tactgaaaaa aatggtccgg cccatctggg
     3301 ctaatgtatt taactctgca ctggcaacaa taacagcagc cattatctac ctgtaccctt
     3361 ctccactggt gtgcaatgcc ggagcctttg ttatcggttt ctcagcagct ggcggcattt
     3421 tacagctcgg cgtttcggtc atgtcagagt tttttcccaa aagcaaagcc aaagtcacca
     3481 gtatttatat gatgatgggt ggactggcta actttgttat tccactgatt accggttatc
     3541 tgtcgaacat cggcctgcaa tatatcattg ttctcgattt tactttcgcg ctgctggccc
     3601 tgattaccgc aattattgtt tttatccgct attaccgcgt tttcattatt cctgaaaatg
     3661 atgtgcggtt tggcgagcgt aaattttgca cccggttaaa cacaattaag catagaggtt
     3721 aa
//
```

**sJEC054**

```
LOCUS       sJEC054          3583 bp ds-DNA     linear
```

```
FEATURES             Location/Qualifiers
     gene            complement(249..1610)
                     /label="ffh"
     CDS             complement(249..1610)

/translation="MFDNLTDRLSRTLRNISGRGRLTEDNVKDTLREVRMALLEADVALPVVREFINRVKEKAVGH
EVNKSLTPGQEFVKIVRNELVAAMGEENQTLNLAAQPPAVVLMAGLQGAGKTTSVGKLGKFLREKHKKKVLVVSAD
VYRPAAIKQLETLAEQVGVDFFPSDVGQKPVDIVNAALKEAKLKFYDVLLVDTAGRLHVDEAMMDEIKQVHASINP
VETLFVVDAMTGQDAANTAKAFNEALPLTGVVLTKVDGDARGGAALSIRHITGKPIKFLGVGEKTEALEPFHPDRI
ASRILGMGDVLSLIEDIESKVDRAQAEKLASKLKKGDGFDLNDFLEQLRQMKNMGGMASLMGKLPGMGQIPDNVKS
QMDDKVLVRMEAIINSMTMKERAKPEIIKGSRKRRIAAGCGMQVQDVNRLLKQFDDMQRMMKKMKKGGMAKMMRSM
KGMMPPGFPGR*"
     misc_feature    1726..1730
                     /label="Transposon site duplication"
     misc_feature    1731..1738
                     /label="Transposon site duplication"
     misc_feature    1731..1875
                     /label="Transposon left end"
     CDS             complement(1912..2091)
                     /label="ChlR    (Chloramphenicol    acetyltransferase)
(partial)"
     promoter        complement(2092..2126)
                     /label="J23119 promoter"
     CDS             complement(2127..2606)
                     /label="ChlR    (Chloramphenicol    acetyltransferase)
(partial)"
     misc_feature    complement(2630..2740)
                     /label="Transposon right end"
     misc_feature    complement(2733..2740)
                     /label="Transposon site duplication"
     misc_feature    2741..2745
                     /label="Transposon site duplication"
     misc_feature    complement(2791..2822)
                     /label="ffh crRNA 1"
     gene            2792..3583
                     /label="ypjD"
     CDS             2792..3583

/translation="MPVFALLALVAYSVSLALIVPGLLQKNGGWRRMAIISAVIALVCHAIALEARILPDGDSGQN
LSLLNVGSLVSLMICTVMTIVASRNRGWLLLPIVYAFALINLALATFMPNEYITHLEATPGMLVHIGLSLFSYATL
IIAALYALQLAWIDYQLKNKKLAFNQEMPPLMSIERKMFHITQIGVVLLTLTLCTGLFYMHNLFSMENIDKAVLSI
VAWFVYIVLLWGHYHEGWRGRRVVWFNVAGAVILTLAYFGSRIVQQLIS*"
ORIGIN
        1 aacatcctct tgtgtgaata aaacaaccgg accccatcga ggaacggagt ccggtgtcat
       61 attaaaagcc cgaaaatttt actcattttt gcgggaattg caatcaacag ttgctaactc
      121 tgctgtaaaa ggccgtcggc ggtgcagcca gtttggtgcc ggagtgcgcg cagtcaccgg
      181 agcgtacacg cagtacgtga ggatgacgag cacatcccgg tgccaaaatg gcaaacaagc
      241 caggccgatt agcgaccagg gaagcctggg ggcatcatac ccttcatgct tctcatcatc
      301 ttcgccattc cgcccttctt cattttcttc atcatgcgct gcatgtcgtc gaactgtttc
      361 agaagacggt taacgtcctg cacctgcata ccgcaaccgg cagcaatacg gcgtttacgc
      421 gaacctttga tgatttctgg cttagcgcgc tctttcatcg tcatcgagtt gatgatggct
      481 tccatacgca ccagcacttt atcgtccatc tgtgacttga cgttatccgg gatctgcccc
      541 atgcccggca gcttgcccat cagactagcc atgccgccca tatttttcat ctggcgcagc
      601 tgctcaagaa agtcgttgag atcgaagccg tcaccttttt tcagcttgct ggctaatttc
      661 tctgcctgcg cgcggtcaac tttgctttcg atatcttcga tcagcgacag tacgtcgccc
      721 atgccgagaa tacgcgacgc gatgcggtcc ggatggaacg gctccagcgc ctcagttttc
      781 tcgccaacac cgaggaactt gatcggtttg ccagtgatgt gacgaataga gagcgccgca
```

```
     841 ccgccgcggg catcgccgtc cactttggtc aacactacgc cggtaagcgg taacgcttca
     901 ttgaatgctt ttgccgtatt ggccgcatcc tgaccggtca tggcgtcaac cacaaacagg
     961 gtttcaaccg ggttaatcga cgcatggact tgtttgatct cgtccatcat cgcttcgtca
    1021 acgtgcagac gaccagcggt atccaccagc agcacgtcgt agaatttcag tttggcttct
    1081 ttcagcgccg cgttaacgat atctaccggc ttctgaccaa catcagaagg gaagaaatca
    1141 acgcccacct gctctgccag cgtctcaagc tgtttgattg ccgccgggcg ataaacgtcg
    1201 gcagaaacca ccagcacttt cttcttgtgc ttctcgcgca ggaacttacc gagtttacca
    1261 acgctggttg ttttaccggc accttgcagg cccgccatca gtacgaccgc aggcggttgc
    1321 gcagccaggt tcagggtctg gttctcttcg cccatcgccg caaccagttc gttacggact
    1381 attttgacga actcctgccc cggcgtcagg ctcttattaa cttcatgacc aaccgctttc
    1441 tcttttacgc gattgataaa ctcacgcact accggcagag ctacgtccgc ctccagcagc
    1501 gccatgcgca cttcgcgcag cgtatctttt acgttgtctt cagtgaggcg tccacggcca
    1561 ctgatattgc gcagcgtgcg cgacaaacga tcggttaaat tatcaaacat tgtctctcgc
    1621 ctggggtgga aacggttggc cgcaatcgcg acacatcatc agtattttgc cgcagtataa
    1681 catgaaggcg tctttgttgt tatgcaacgg ttggagcagc gttcacctga tgttgatgca
    1741 accataaagt gatatttaat aattatttat aatcagcaac ttaaccacaa aacaaccata
    1801 tattgatatc tcacaaaaca accataagtt gatatttttg tgaatcgagt atttcagcaa
    1861 aactactgca gtaaggactc tagaagggca ccaataactg ccttaaaaaa attacgcccc
    1921 gccctgccac tcatcgcagt actgttgtaa ttcattaagc attctgccga catggaagcc
    1981 atcacaaacg gcatgatgaa cctgaatcgc cagcggcatc agcaccttgt cgccttgcgt
    2041 ataatatttg cccatagtga aaacgggggc gaagaagttg tccatattgg cgctagtatt
    2101 atacctagga ctgagctagc tgtcaacacg tttaaatcaa aactggtgaa actcacccag
    2161 ggattggctg agacgaaaaa catattctca ataaaccctt tagggaaata ggccaggttt
    2221 tcaccgtaac acgccacatc ttgcgaatat atgtgtagaa actgccggaa atcgtcgtgg
    2281 tattcactcc agagcgatga aaacgtttca gtttgctcat ggaaaacggt gtaacaaggg
    2341 tgaacactat cccatatcac cagctcaccg tctttcattg ccatacggaa ctccggatga
    2401 gcattcatca ggcgggcaag aatgtgaata aaggccggat aaaacttgtg cttatttttc
    2461 tttacggtct ttaaaaaggc cgtaatatcc agctgaacgg tctggttata ggtacattga
    2521 gcaactgact gaaatgcctc aaaatgttct ttacgatgcc attgggatat atcaacggtg
    2581 gtatatccag tgattttttt ctccatggat ccccggaaat gacctcgagg aaaatgtcgt
    2641 aaacttactg aagtttagac catgaagagg caatatcaat ttatgggtgt gataattatc
    2701 aatttatggg tgtaattatc attttatggt tgtatcaaca cctgacgcta tactgcttct
    2761 ctttcttatt gctcaaactg tcgacatcac tatgcccgtt tttgctctgc tcgcgcttgt
    2821 cgcctactcc gtcagtcttg cgctgattgt tcccggtctg ctgcaaaaaa acggcggctg
    2881 gcggcgcatg gctattattt ctgcggtcat tgcgctggtc tgccacgcaa tcgctctgga
    2941 agcccgcatc ctgcccgacg gtgatagcgg acaaaacctc agcctgctga acgttggttc
    3001 attggtcagt ttgatgatct gtacggtaat gaccattgtg gcttctcgca atcgtggctg
    3061 gctgctgcta cccattgtct atgcctttgc gcttatcaac ctggcgctgg caaccttcat
    3121 gcccaatgaa tacatcaccc atctggaagc tacgcctggg atgctggtgc acattggctt
    3181 atcgctcttt tcctatgcca cgctaattat cgccgccctg tacgcgctgc aactggcgtg
    3241 gattgattac caactgaaga acaagaagct ggcgtttaac caggaaatgc cgccattgat
    3301 gagtatcgag cgtaaaatgt tccacatcac gcagattggc gtggtgctgc taacgctcac
    3361 gctttgcact ggcctgttct acatgcacaa cttatttagc atggaaaata tcgacaaggc
    3421 tgtgctctct atcgtggcgt ggtttgtcta tattgtgctg ctgtggggac attatcatga
    3481 aggatggcgt ggacgccgcg tcgtctggtt taacgttgcg ggcgcggtaa ttctgacact
    3541 ggcctacttc ggcagccgaa ttgtccagca gttaatcagc taa
//
```

| Supplementary Table 4. crRNAs used in this study. | |
|---|---|
| **crRNA ID** | **Sequence (5' to 3')** |
| mRFP1 | ATTGTTAGCCGTATGAATAGGGAAACTAAACC |
| non-targeting | GAGACCTCTGTCTTGTCAGCTAGGGTGGTCTC |
| cadB | TCAAGTTCTCACTTACAGAAACTTTTGTGTTA |
| ffh | CGACAAGCGCGAGCAGAGCAAAAACGGGCATA |
| ydiN | AAGGAAAATCGCGATCAAAAACAAACTATGAC |
| zraP | TGCGCTGTAACACAAAGGCTTAAGTTTCAATG |
| ampC 1 | GTTGTCACGCTGATTGGTGTCGTTACAATCTA |
| ampC 2 | TGACAGTTGTCACGCTGATTGGTGTCGTTACA |
| marA 1 | AAGAATTAACAAAAAACCTGACGGCGGACGAA |
| marA 2 | GCACCAAGAATTAACAAAAAACCTGACGGCGG |

**Supplementary Table 5. Primers used in this study.**

| ID | Sequence (5' to 3') | Used for figures |
|---|---|---|
| NGS fwd 1 | ACACTCTTTCCCTACACGACGCTCTTCCGATCTACGTCCCAATATATGGTTGTTTTGTGGTTAAGTTGC | SF2B |
| NGS fwd 2 | ACACTCTTTCCCTACACGACGCTCTTCCGATCTTGACAGCAATATATGGTTGTTTTGTGGTTAAGTTGC | |
| NGS fwd 3 | ACACTCTTTCCCTACACGACGCTCTTCCGATCTCTTGGACAATATATGGTTGTTTTGTGGTTAAGTTGC | |
| NGS rev 1 | GACTGGAGTTCAGACGTGTGCTCTTCCGATCTACGTCCGAACCTTCCATACGAACTTTGAAACG | |
| NGS rev 2 | GACTGGAGTTCAGACGTGTGCTCTTCCGATCTTGACAGGAACCTTCCATACGAACTTTGAAACG | |
| NGS rev 3 | GACTGGAGTTCAGACGTGTGCTCTTCCGATCTCTTGGAGAACCTTCCATACGAACTTTGAAACG | |
| cadB qPCR primer 1 | TACCAACATCATCCAGGAGC | 3B |
| cadB qPCR primer 2 | AGTGCTTCAACTATCCTCGG | |
| cadB RT primer | TTCATCAGAGTGATAAGG | |
| ffh qPCR primer 1 | CAAGAAGAAAGTGCTGGTGG | |
| ffh qPCR primer 2 | TCTGACCAACATCAGAAGGG | |
| ffh RT primer | TCGTAGAATTTCAGTTTG | |
| ydiN qPCR primer 1 | GCGGTCATACTGGTTAAAGC | |
| ydiN qPCR primer 2 | ATACCCGGAATAATCAGCCC | |
| ydiN RT primer | TAATTCATTAGTTACGCTG | |
| zraP qPCR primer 1 | ATACAATGCCCTGTTAGCCG | |
| zraP qPCR primer 2 | ACTCATCTAACGACTGACGC | |
| zraP RT primer | CATACCCATATGACCG | |
| cadB fwd | AACTCCGGGTTGATTTATGCTC | SF6 |
| cadB rev | ATAATGCAATACCGCTCCCC | |
| ffh fwd | GCGCAAGACTGACGGAGTAG | |
| ffh rev | TCAGGGTCTGGTTCTCTTCG | |
| ydiN rev | TACTGATCAATCGCCCCAAAC | |
| ydiN fwd | GCACTGTTTATCGGCTTACG | |
| zraP fwd | AGCGCGGATAAGCACATTAC | |
| zraP rev | CAGGGCAATTTTCGTGTTCC | |
| marA rev | TTGCGACTCGAAGCCATATC | |
| marA fwd | CTGTAAAGGCTGGGTGGAAAG | |
| ampC fwd | GGCTGCTATCCTGACAGTTG | |
| ampC rev | TTGATTTGTTGAGGGGCAGC | |
| Tn RE | CTCGAGGAAAATGTCGTAAACTTACTG | |
| Tn LE | CAATATATGGTTGTTTTGTGGTTAAGTTGC | |
